# Circadian period is compensated for repressor protein turnover rates in single cells

**DOI:** 10.1101/2024.02.06.579141

**Authors:** Christian H. Gabriel, Marta del Olmo, Arunya Rizki Widini, Rashin Roshanbin, Jonas Woyde, Ebrahim Hamza, Nica-Nicoleta Gutu, Amin Zehtabian, Helge Ewers, Adrian Granada, Hanspeter Herzel, Achim Kramer

## Abstract

Most mammalian cells possess molecular circadian clocks generating widespread rhythms, e.g. in transcript and protein abundance. While circadian clocks are robust to fluctuations in the cellular environment, little is known about how circadian period is compensated for fluctuating metabolic states. Here, we exploit the heterogeneity of single cells both in circadian period and metabolic state, governing protein stability, to study their interdependence without the need for genetic manipulation. We generated cells expressing key circadian proteins (CRY1/2 and PER1/2) as endogenous fusions with fluorescent proteins and simultaneously monitored circadian rhythms and degradation in thousands of single cells. We found that the circadian period is compensated for fluctuations in the turnover rates of circadian repressor proteins and uncovered possible mechanisms using a mathematical model. In addition, the stabilities of the repressor proteins are circadian phase-dependent and correlate with the circadian period in a phase-dependent manner, in contrast to the prevailing model.

## Introduction

Circadian clocks have evolved in all kingdoms of life, enabling organisms to track, anticipate and adapt to the ∼24-hour rhythm of day and night. They exist at all levels of hierarchy, from single cells to organs and whole organisms - but the basis of all circadian rhythms is a cell-autonomous oscillator.^1^ However, the dynamics of single-cell circadian rhythms have a high degree of noise and stochasticity, e.g. the circadian clock of individual cells can oscillate with periods ranging from about 18 hours to 30 hours and beyond, despite being genotypically identical.^2–5^ Intercellular communication allows noisy single-cell oscillations to give rise to a robust rhythmic signal at the population or organ level.^6,7^ In mammals, the generation of molecular oscillations is thought to rely on a transcriptional-translational feedback loop (TTFL): CLOCK:BMAL1 promotes the rhythmic expression of the repressors CRY1-2 and PER1-3, by binding to E-box elements in their promoters.^8^ PERs and CRYs form high molecular-weight complexes that inhibit CLOCK:BMAL1, thereby repressing their own transcription.^9–12^ After the regulated degradation of the complex, the monomeric CRY1 still independently represses CLOCK:BMAL1 until the inhibition is released and a new cycle begins.^13,14^

A fundamental question in circadian biology remains: what determines the circadian period length? The mere structure of the TTFL cannot explain this and critical delays are required to enable oscillation in the first place and extend the duration of this loop to ∼24 h.^15^ After Konopka and Benzer discovered that phosphorylation mutants of the *Drosophila period* gene cause period phenotypes, it became clear that post-translational modifications (PTMs) play important roles in period determination^16,17^, e.g. by modulating the stability of the repressors. In fact, mutation, inhibition or ablation of ubiquitin ligases targeting CRYs and PERs for degradation, e.g. FBXL3 and βTrCP, not only increases the protein half-life, but also lengthens the period^18–24^, suggesting that repressor stability and circadian period may be directly correlated. However, this concept did not hold when circadian oscillations were restored by introducing CRY1 into arrhythmic *Cry*1/2-knockout cells^13^: although for several CRY1 mutants the rescued period did indeed correlate with protein stability, other mutants did not fit this pattern.^25–27^ Genetic manipulation by mutation almost always carries the risk of also altering protein function, and thus a period phenotype may occur because of, in addition to, or despite the alteration in protein stability. Therefore, while mutants can be valuable tools, they also have clear limitations.^25,28^

Here we exploit the natural heterogeneity of both the circadian period and protein degradation rates at the single cell level, which allows us to study the interdependence of these traits without the need for genetic manipulation.^2,4,29^ Using tens of thousands of engineered single cells expressing CRYs and PERs as fusion proteins with fluorescent reporters, we found that the stability of these proteins is far from constant, but varies with the time of day, which, in addition to circadian transcription, conditions rhythmic protein levels. The influence of repressor stability on the circadian period also turned out to be phase-dependent: early in the cycle, high stability correlates with a shorter period, late in the cycle with a longer period. Overall, however, the circadian period is surprisingly resilient to strongly fluctuating protein degradation rates. We reproduce and conceptualize these findings with a mathematical model that describes several interacting mechanisms for this compensation of the circadian period against highly variable protein degradation rates.

## Results

### Visualization of endogenous CRY2 and PER1 proteins in living cells

We have previously used CRISPR/Cas9-mediated knock-in approaches to generate U-2 OS cells expressing CRY1 and/or PER2 as fluorescent fusion proteins from the endogenous locus^4^. In these cells, the nuclear accumulation of both fusion proteins oscillates in a circadian manner. Because the paralogues PER1 and CRY2 have overlapping but not redundant functions within the TTFL, ^30–33^ and to study the protein dynamics of all circadian repressors side by side in living cells, we generated CRY2-mScarlet-I and PER1-mScarlet-I knock-in cells (referred to as CRY2-mSca and PER1-mSca, respectively). We used a similar Cas9-mediated HDR approach to insert the sequence of the red fluorescent protein mScarlet-I (mSca) 5’ to the PER1 or CRY2 stop codon into the genome of U-2 OS cells (**Supplementary Fig. S1A**) and screened clones by fluorescence microscopy and genomic PCR. We selected two homozygous CRY2-mSca-I and two heterozygous PER1-mSca clones (**Supplementary Fig. S1B-E**) and confirmed the specificity of the fluorescence using shRNA targeting CRY2 and PER1, respectively (**Fig. 1A-B**). Circadian rhythms were intact in all clones, as demonstrated by rhythmic activation of a Bmal1::Luc reporter with similar circadian dynamics compared to wild-type cells (**Supplementary Fig. S1F-J**). CRY2-mSca fluorescence was seen exclusively in the nucleus of the knock-in cells (**Fig. 1A**), similar to what we have observed for CRY1^1^. While fluorescence signals in PER1 knock-in cells were mainly detected in the nucleus, fluorescence levels above background were also observed in the cytoplasm (**Fig. 1B**). Taken together, these results indicate successful expression of the fluorescent CRY2-mSca or PER1-mSca fusion protein from the endogenous genomic loci in these clones, while the circadian oscillator remains intact.

**Figure 1:**
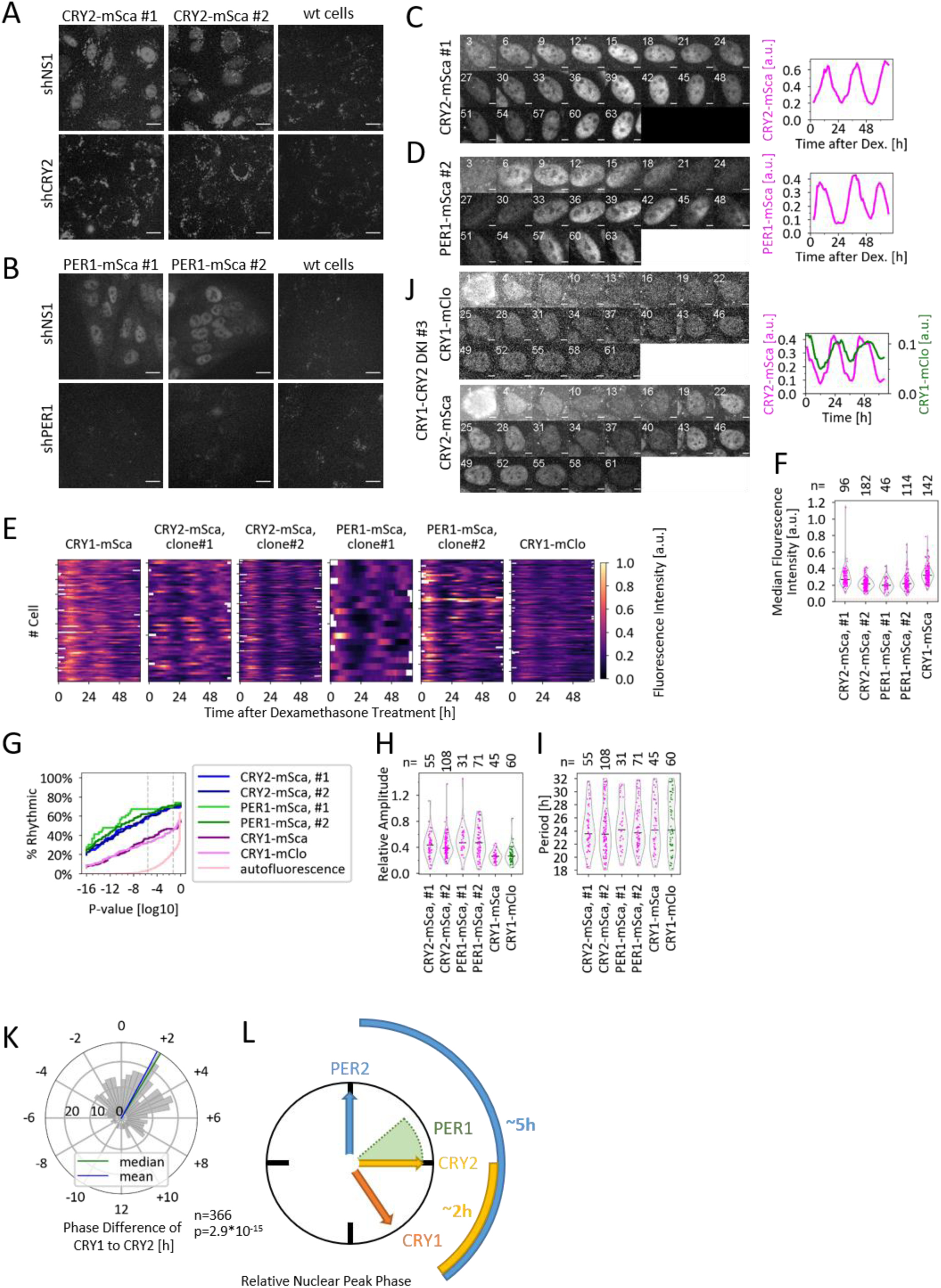
CRY2 and PER1 protein oscillation in single cells. (A,B) Fluorescent knock-in and wild-type (wt) U-2 OS cells transduced with shRNA targeting either *CRY2* or *PER1*, or with a non-silencing control shRNA (shNS1). Scale bar: 20 µm. (C,D) Montage of a CRY2-mSca (C) or PER1-mSca (D) knock-in cell nucleus recorded at indicated hours after dexamethasone (dex) treatment, and time course of mean fluorescence intensity. Scale bar: 5 µm.(E) Time series of mean nuclear fluorescence of indicated knock-in clones after dex treatment. Only time series with ≥ 60 h are shown. y-axis ticks mark every 10^th^ cell. (F) Median fluorescence intensities for individual cells from all time points after background subtraction. Horizontal lines: median of all cells, red line: median nuclear autofluorescence. (G) Percentage of rhythmic cells as a function of p-value cutoff. Vertical dashed lines represent p-values of 0.05 and the more stringent value used here, respectively. See **Supplementary Note 1.** (H,I) Relative amplitudes (H) and periods (I) of rhythmic time series. (J) Montage of a CRY1-mClo/CRY2-mSca double knock-in (DKI) cell nucleus imaged in the two fluorescence channels at the indicated hours after dex treatment, and time course of mean fluorescence intensities. Scale bar: 5 µm. (K) Histogram of phase difference (30-min bins) in unsynchronized CRY1/CRY2 double knock-in cells. p-value: Wilcoxon signed-rank test. (L) Sequence of mean nuclear peak expression in U-2 OS cells. PER1 timing is estimated from **Supplementary Fig. S3B-C.**

### CRY2 and PER1 protein abundance oscillates in single cells

The protein abundance of both PER1 and CRY2 is known to oscillate over the course of a day at the population level,^34,35^ but little is known about their expression dynamics in single cells. To address this, we recorded fluorescence of single cells from our newly generated knock-in clones for three days after dexamethasone synchronization with a time resolution of 1h. We observed that the nuclear abundance of both PER1-mSca and CRY2-mSca oscillated in single cells, but with different characteristics. CRY2-mSca was detected in the nucleus of expressing cells throughout the circadian cycle and did not exceed background levels in the cytoplasm at any time point (**Fig. 1C**). In contrast, nuclear PER1-mSca levels of many cells dropped to near background fluorescence levels in the trough of the circadian oscillation (**Fig. 1D**).

To quantify circadian dynamics, we developed an automated approach to track and extract signals from thousands of cells in a single experiment. Briefly, cells were stably transduced to express a nuclear infrared protein (histone-2B-miRFP720). After imaging, nuclei were segmented and tracked based on miRFP720 fluorescence using Cellprofiler software, and the background-subtracted mean nuclear fluorescence from different channels was extracted. To improve data quality, mistracked nuclei were identified by an apparent abrupt size change in the absence of cell division and filtered out using a Python script (see Methods). Using this approach, we extracted nuclear fluorescence signals from hundreds of dexamethasone-synchronized PER1-mSca, CRY2-mSca, and - as a reference - CRY1-mSca and CRY1-mClover3 reporter cells over the course of 68 hours (**Fig. 1E**, **Supplementary Fig. S2, Supplementary Video SV1**). In contrast to what we have observed for PER2 - which was expressed 6-10 times lower than CRY1^4^ - the average intensity levels of PER1-mSca, CRY1-mSca and CRY2-mSca were similar (**Fig. 1F**).

Next, we analyzed the circadian rhythmicity of these time series using Metaycycle2D, which integrates nonparametric and Lomb-Scargle periodogram analysis^36–38^ to calculate circadian parameters and a p-value for rhythmicity (**Supplementary Fig. S2**). We defined a dataset-specific stringent p-value cut-off based on the autofluorescence recording (p<10^-5^, see **Supplementary Note 1** for details) and excluded time series that fell below this value or whose calculated period exactly matched the entered limits (18-32 h), as the latter are most likely to contain oscillations longer or shorter than these limits. Using these criteria, 59-66% of the time series from CRY2-mSca and PER1-mSca cells were classified as ‘highly rhythmic’, compared to ∼33% of those from CRY1 reporter cells. These differences between repressor oscillations were present regardless of the p-value cutoffs (**Fig. 1G**). Among the highly rhythmic time series, PER1 protein oscillations had the highest relative amplitude, while CRY1 oscillations had the lowest (**Fig. 1H**). The average periods of the rhythmic signals were similar for all six clones analyzed (**Fig. 1I**). Notably, the periods of individual cells - even within clonal populations - were highly variable, covering the full range of 18-32 hours.

### Phase relationship between CRY and PER proteins

Previously, we observed that the expression phase of CRY1 protein was delayed by approximately 5 h relative to that of PER2 in dual reporter cells^4^, which is consistent with a delayed mRNA expression of CRY1 relative to the other circadian repressor proteins and ChIP-Seq time series showing exclusive presence of CRY1 at E-boxes in a late repression phase^14,35^.

To obtain a more complete and time-resolved profile of circadian repressor expression, we also aimed to estimate the expression phase of CRY2 and PER1 in relation to CRY1. Overall, our rhythmic cells had a low phase coherence of clonal cell populations, i.e., the circadian phases at 2 and 3 days after synchronization were quite different (**Supplementary Fig. S3A**). While this was consistent with the single-cell heterogeneity observed for circadian periods, it made it difficult to calculate a significant average peak phase. To estimate the average phase at the population level, we calculated the mean of all normalized time series and determined the time of the second peak after synchronization (**Supplementary Fig. S3B-C)**). This revealed that the nuclear accumulation of PER1 peaked first, followed by CRY2 and CRY1 last, with the limitation that the clonal difference was larger than the differences between the reporters.

To overcome this limitation, we generated double knock-in cells expressing CRY1-mClover together with either PER1-mSca or CRY2-mSca. These double knock-in cells allowed us to visualize and study the dynamics of different repressor proteins in parallel within the same cell. Unfortunately, putative PER1-mSca/CRY1-mClo double knock-in cells unexpectedly showed an exclusively cytoplasmic localization of the mSca fluorescence signal, suggesting a deleterious interplay of the two fusion proteins. However, we successfully generated CRY1- mClover3/CRY2-mSca double knock-in cells in which both mClo and mSca fluorescence signals were localized to the nucleus, as seen in cells expressing either fusion protein alone (**Fig. 1J**, **Supplementary Fig. S4A**). Knock-in was verified by genomic PCR (**Supplementary Fig. S4B-C**) and the specificity of the fluorescence signal was confirmed by shRNA-mediated knockdown (**Supplementary Fig. S4A**) for four clones. These clones showed similar circadian rhythmicity compared to their respective parental clone (**Supplementary Fig. S4D-H**) allowing us to simultaneously monitor both CRY proteins in single cells with intact circadian clocks (**Fig. 1J**).

We monitored the nuclear fluorescence of unsynchronized CRY1/CRY2 double knock-in cells over the course of two days and identified cells in which the signals of both proteins were rhythmic. In these cells, CRY2 nuclear accumulation peaked on average 1.9±4.3 hours (mean ± SD) before that of CRY1 (**Fig. 1K**). The high standard deviation again demonstrated the high degree of variability of protein oscillations in single cells. Thus, from our data, we propose the following sequence of events in the nucleus of U-2 OS cells: (1) peak of PER2 protein, (2) peak of PER1 protein ∼1.5-3 hours later, (3) peak of CRY2 protein 3 hours after the peak of PER2 and finally peak of CRY1 protein another 2 hours later (**Fig. 1L**).

### Stability of repressor proteins

Using these circadian reporter cells, we sought to address the fundamental unresolved question of how the period of the molecular circadian clock is tuned to ∼24 hours. While there is evidence that altering the stability of circadian repressor proteins, i.e., CRYs and PERs, can also affect the circadian period,^18,23^ such data have mostly been derived from genetic perturbation studies, and it remains unclear whether the altered period is a consequence of altered stability or of the perturbation itself. Given the high variability of circadian oscillations in single cells, we hypothesized that we could exploit the heterogeneity of single cells to analyze the interdependence of repressor stability and circadian period without the need for genetic manipulation. To simultaneously obtain circadian parameters and repressor protein stability in the same cells, we monitored nuclear protein abundance of unsynchronized single and double knock-in cells (**Supplementary Tab. S1)** over the course of three days, capturing rhythms in the first two days and recording degradation dynamics on the third day by then stopping any new protein synthesis with cycloheximide (CHX, 20 µg/ml, **Supplementary Video SV2**). Circadian parameters and the circadian phase at which CHX was added – and at which protein half-life is assessed - were calculated from time points prior to CHX addition using Metacycle2D. Protein half-lives were obtained by fitting mono-exponential decay curves to the time series after CHX addition (**Fig. 2A**).

**Figure 2:**
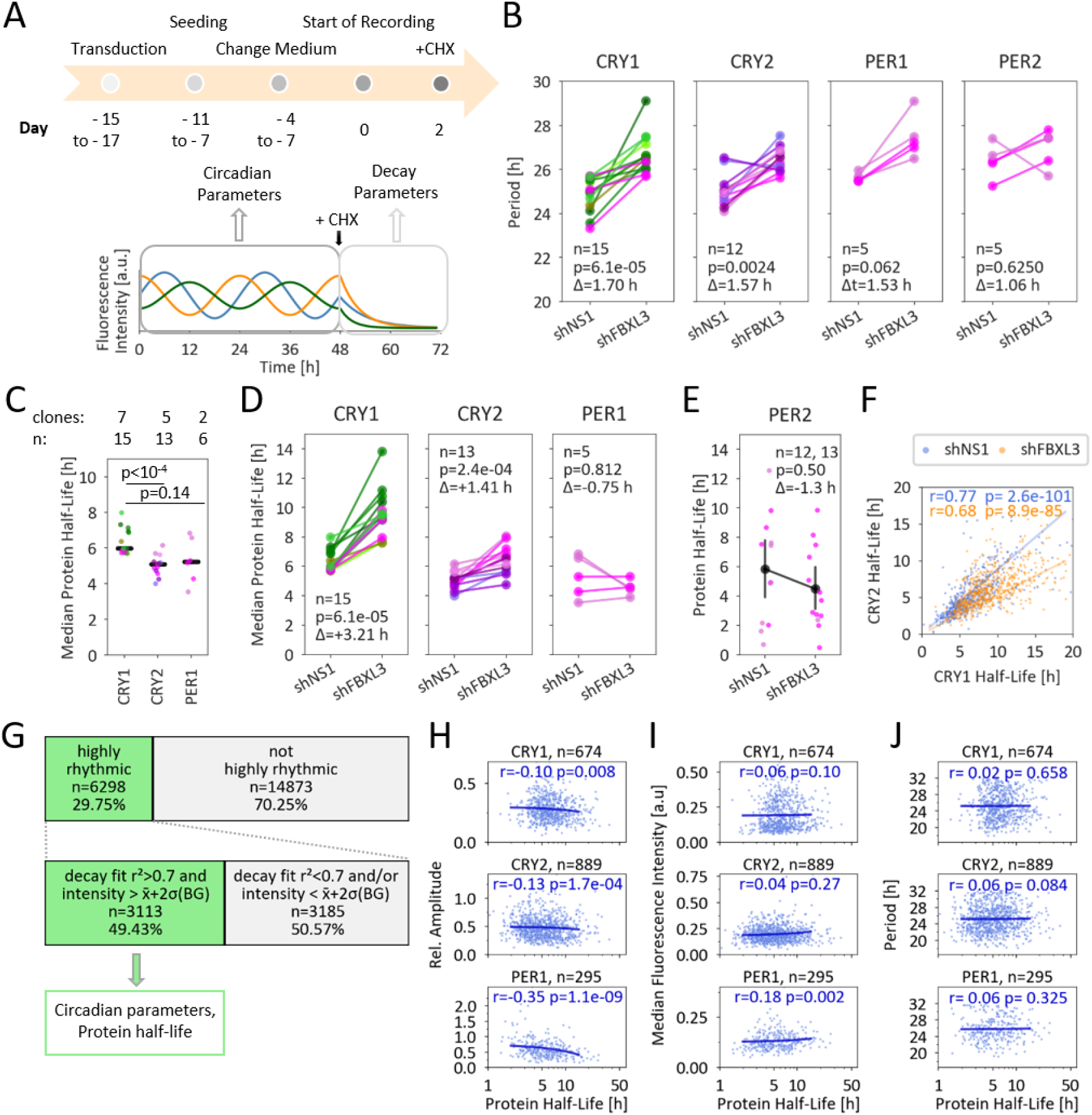
Stability of repressor proteins and correlation with circadian dynamics. (A) Experimental setup: Unsynchronized single knock-in reporter cells transduced with shRNA were imaged 2 days prior and 1 day after cycloheximide addition. Circadian parameters were extracted from days 1 and 2, and protein half-life from day 3.(B) Median periods of reporter cells transduced with the indicated shRNAs, each calculated from ≥10 rhythmic cells. Same colors (green: mClo clones, purple: mSca clones) represent the same clonal population from different experiments. p-value: Wilcoxon signed-rank test. (C,D) Median protein half-life of clonal populations transduced with control shRNA (C) or the indicated shRNAs (D), each calculated from ≥10 decay fits. Black lines (C) indicate median. p-values: Mann-Whitney-U test (C), Wilcoxon signed-rank test (D). (E) Half-life of individual PER2-mSca cells. Shown are means ±95% confidence interval (CI). p-value: Wilcoxon signed-rank test. (F) Correlation of CRY1 and CRY2 half-life in double knock-in cells (Spearman). (G) Highly rhythmic time series (**Supplementary Note 1**) were filtered for reliable decay fits. (H,I,J) Correlation of measured half-lives with relative circadian amplitude (H), signal intensity (abundance, I) and circadian period length (J). Correlation coefficient (r) and p-value from Spearman correlation, line: linear regression.

To test our ability to accurately determine periods and protein half-lives in single cells, we sought to reproduce the observation from population studies that knockdown of the ubiquitin ligase FBXL3 results in a long circadian period and increased CRY half-lives^23^. To this end, cells were transduced with either a non-silencing shRNA or an shRNA targeting FBXL3. In total, we obtained more than 20,000 time series of nuclear fluorescence from three independent experiments (**Supplementary Tab. S2**). Using the same threshold criterion as described above, approximately 30% of all time series were classified as highly rhythmic (24.4% for FBXL3 knockdown and 36.9% for control cells, **Supplementary Tab. S2**). Cells from most clones oscillated with average periods between 23 and 25 hours and had various phases when CHX was added (**Supplementary Fig. S5A**). Knockdown of FBXL3 increased the average period of almost all clonal populations by 1.5±1.1 hours (mean ± SD, **Fig. 2B**), similar to what has been described previously^23^. This demonstrates that despite the high cell-to-cell variability of circadian periods, differences in period distribution can be faithfully detected in single cell data. Next, we calculated the half-life of the mean nuclear fluorescence signal after addition of CHX, which we will refer to as the half-life of the respective protein for ease of reading. After correction for photobleaching (see Methods), we fitted mono-exponential decay curves to the second part of the time series, starting 2 hours after addition of CHX. We required (i) that the initial intensity be significantly above background levels (∼54% of all traces), and (ii) a r² value of at least 0.7 for a successful fit (9211 traces, ∼82%).

Overall, the protein half-lives obtained were highly variable, covering almost an order of magnitude, with 95% of the values falling between 2.3 and 19.0 hours (**Supplementary Fig. S5B-C**). Comparing the median half-lives of the repressor proteins of the clonal populations, we observed that on average CRY1 proteins had a significantly longer half-life (5.9±0.7 h, median ± SD) than CRY2 (5.1±0.6 h, p=9.3*10^-5^, Mann-Whitney-U test) and, although not statistically significant, PER1 (5.2±1.2 h, p=0.14, **Fig. 2C**). For PER2, we could only reliably determine the half-life in 25 cells (5.8±4.6 h, **Fig. 2E**) because the signal’s intensity was too low in most cells. Therefore, we had to refrain from further statistical analysis of PER2 half-lives and instead focus on the other three repressors. Consistent with previous reports and the established role of FBXL3 as a CRY ubiquitin ligase, FBXL3 knockdown significantly increased the average half-life of CRY1 and CRY2 (by 3.2 h and 1.4 h, p = 6.1*10^-5^ and 1.2*10^-4^, respectively, Wilcoxon signed rank test), but not of PER proteins (**Fig. 2D-E**). Thus, we were able to detect known changes in protein half-lives using noisy single cell data, despite the wide distribution of half-lives within clonal populations (**Supplementary Fig. S5C**). Furthermore, the half-lives of the fusion proteins represent those of the repressors and not those of the fluorescent reporter because, first, only CRY, but not PER fusion protein half-lives increased after FBXL3 knockdown, and second, fluorescent protein half-lives are typically much longer.^39^ Interestingly, we observed a strong correlation between CRY1 and CRY2 half-lives in cells expressing both reporters, which was also evident when FBXL3 was knocked down (**Fig. 2F**). Thus, the stability of CRY1 and CRY2 appears to be co-regulated.

To investigate the effect of repressor stability on circadian dynamics, we focused on those ∼3100 highly rhythmic time series for which we also obtained reliable decay fits (**Fig. 2G**). We observed the expected positive correlation of half-life with expression level (magnitude, **Fig. 2H**) and the negative correlation of half-life with relative amplitude (**Fig. 2I**).^40^ Surprisingly and in contrast to the prevailing model (see above), we did not detect a significant correlation between repressor half-life and circadian period (**Fig. 2J, Supplementary Fig. S5D**). In cells with similar circadian periods, the half-life of e.g. CRY1 can differ by a factor of up to 10 (**Fig. 2J**).

### Protein stability of repressor proteins changes with circadian phase

We speculated that the half-life of CRY1, CRY2, and PER1 may not be a static value, but may itself be subject to circadian changes, as reported for PER2^41^. If this were the case, the measured decay rates would represent a phase-dependent snapshot rather than constant rates, and as such may be insufficient to explain the period of a cell. Indeed, we observed that the half-life of all three proteins showed significant rhythmicity, with a peak of stability during the rising phase (**Fig. 3A**). To get a clearer picture of how the repressor half-life changes during different phases, we binned the half-life and relative expression data into 24 overlapping phase windows of 0.78 rad, corresponding to 3 hours of a 24-hour cycle (**Supplementary Fig. S6**). Thus, by definition, each cell is represented in 3 consecutive bins. Plotting relative expression against circadian phase at the time of CHX addition showed the expected rhythmic patterns, validating the phase determination and binning (**Fig. 3B**). The relationship between half-life and protein expression is as follows (**Fig. 3B-C**): the stability of all three proteins was lowest in cells assayed at or after the peak phase, when protein abundance declines, and highest during and after the trough phase, when proteins reaccumulate. Notably, the number of time series that could be analyzed was lower in the trough phase (**Fig. 3C, lower panels**) because, as with PER2 in general (see above), the low initial signal levels often precluded a faithful determination of the decay dynamics. This is especially true for time series from PER1 reporter cells, whose nuclear trough expression levels are close to background (**Fig. 1D**).

**Figure 3:**
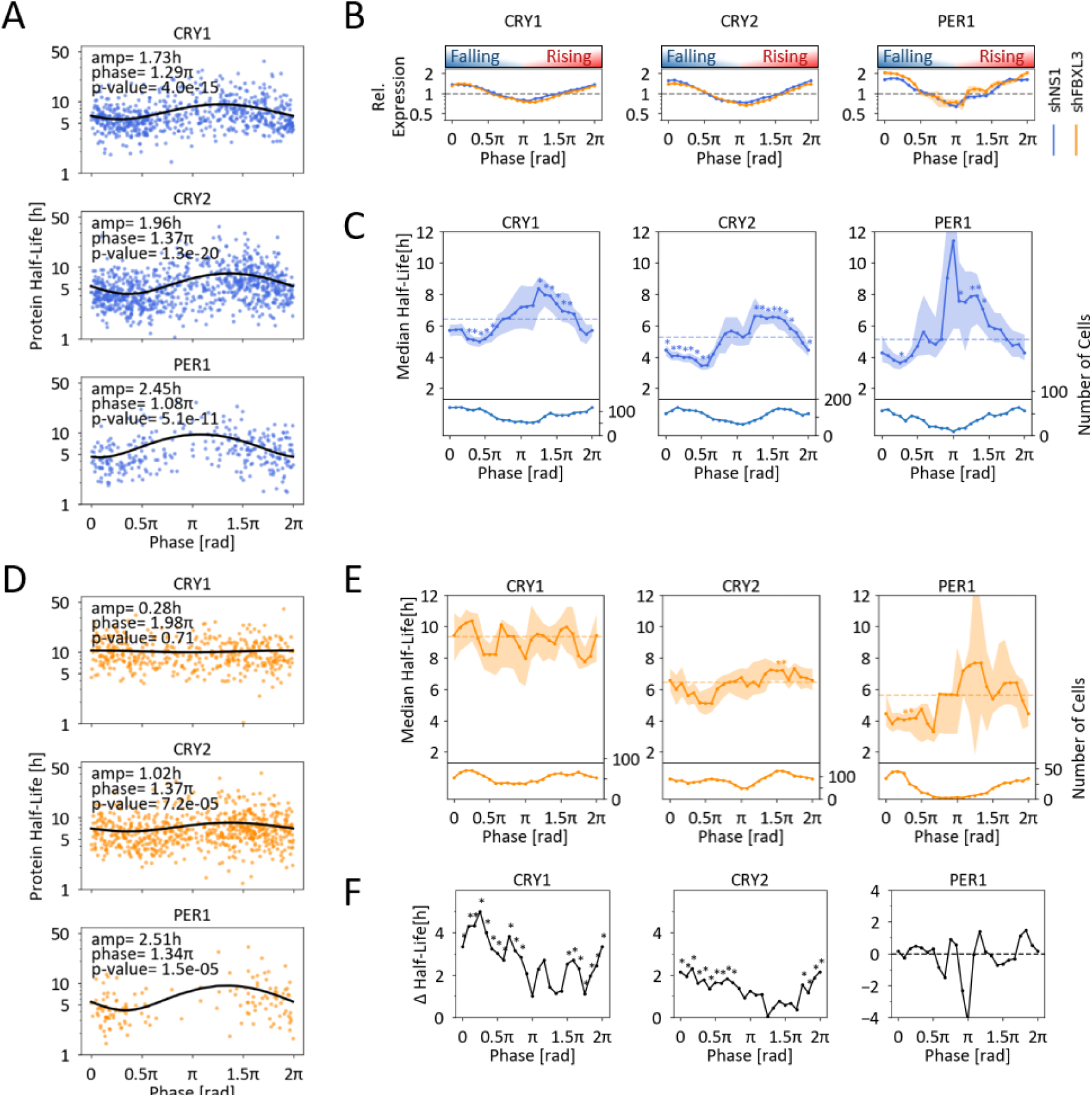
Stability of repressor proteins is circadian phase dependent. (A) Harmonic regression analysis of protein half-life and circadian phase in which the half-life was measured. p-values: F-test. (B) Relative expression at time of CHX addition (mean ± SEM) for each 3-hour phase window. (C) Protein half-life (median and 95% CI) and cell number for each 3-hour phase window. Dashed lines represent median, and * statistically significant (p<0.05) difference from median (1sample Wilcoxon Signed test, corrected for multiple testing (Sidak-Holmes). (D) Harmonic regression analysis of protein half-life and circadian phase after knockdown of FBXL3. p-values: F-test. (E) Protein half-lives as in (C) after knockdown of FBXL3. (F) Increase in median protein half-life in FBXL3 knock-down cells compared to controls, number of cells as in (C) and (E). *: p<0.05, Mann-Whitney-U test, corrected for multiple testing (Sidak-Holmes).

### Rhythmic CRY1 stability is dependent on FBXL3

Next, we asked how the observed rhythmic stabilities might be generated. For the CRY proteins, we speculated that FBXL3 not only affects the average stability (**Fig. 2D**), but acts in a phase-dependent manner. Indeed, we observed that upon FBXL3 knockdown, the observed phase dependence of CRY1 stability is lost (**Fig. 3D**) and CRY1 half-lives are high in all phases (**Fig. 3E**). Thus, the presence of FBXL3 seems to be necessary for the rhythmic stability of CRY1. For CRY2, phase-dependent differences in protein half-life are still present, but are reduced in the absence of FBXL3. In contrast, FBXL3 does not appear to alter the pattern of rhythmic PER1 stability. Interestingly, during the rising phase, when CRY stability was already high, knockdown of FBXL3 did not result in a significant further increase (**Fig. 3F**) suggesting that FBXL3 targets CRY proteins for degradation mainly during the falling phase, resulting in phase- dependent differences in protein stability.

### Impact of CRY half-life on circadian period is phase dependent

Since PER and CRY stabilities change in a phase-dependent manner, the decay rate obtained from a cell represents only a snapshot of a dynamic measure. Therefore, we investigated whether the independence of the period of repressor stability that we found (**Fig. 2J**) could be an artifact of analyzing pooled data. To this end, we grouped time series into overlapping phase windows of 3 hours (as before, **Supplementary Fig. S6B**) and correlated repressor half-life and period separately for cells within these phase windows (**Fig. 4A-E**). Indeed, we found correlations between repressor stability and circadian period, but to an unexpected extent: depending on the circadian phase, we observed either a significantly positive (**Fig. 4B**), negative (**Fig. 4D**) or no correlation (**Fig. 4A,C**) between period and repressor half-life. When the Spearman correlation coefficient and the slope of the linear regression were plotted for all phase windows, a general pattern emerged (**Fig. 4E-H**): in cells assayed during the rising phase, CRY1 stability and circadian period were negatively correlated, i.e. a relatively longer CRY1 half-life in this phase was correlated with shorter periods and vice versa (**Fig. 4B,F**). In contrast, CRY1 half-life and period were positively correlated in cells analyzed during the late falling phase (**Fig. 4D,F**). This was very similar for the phase-dependent correlation of period and CRY2 half-life (**Fig. 4G**). For PER1, the correlation analysis suffered from lower cell numbers and did not show clear trends (**Fig. 4H**).

**Figure 4:**
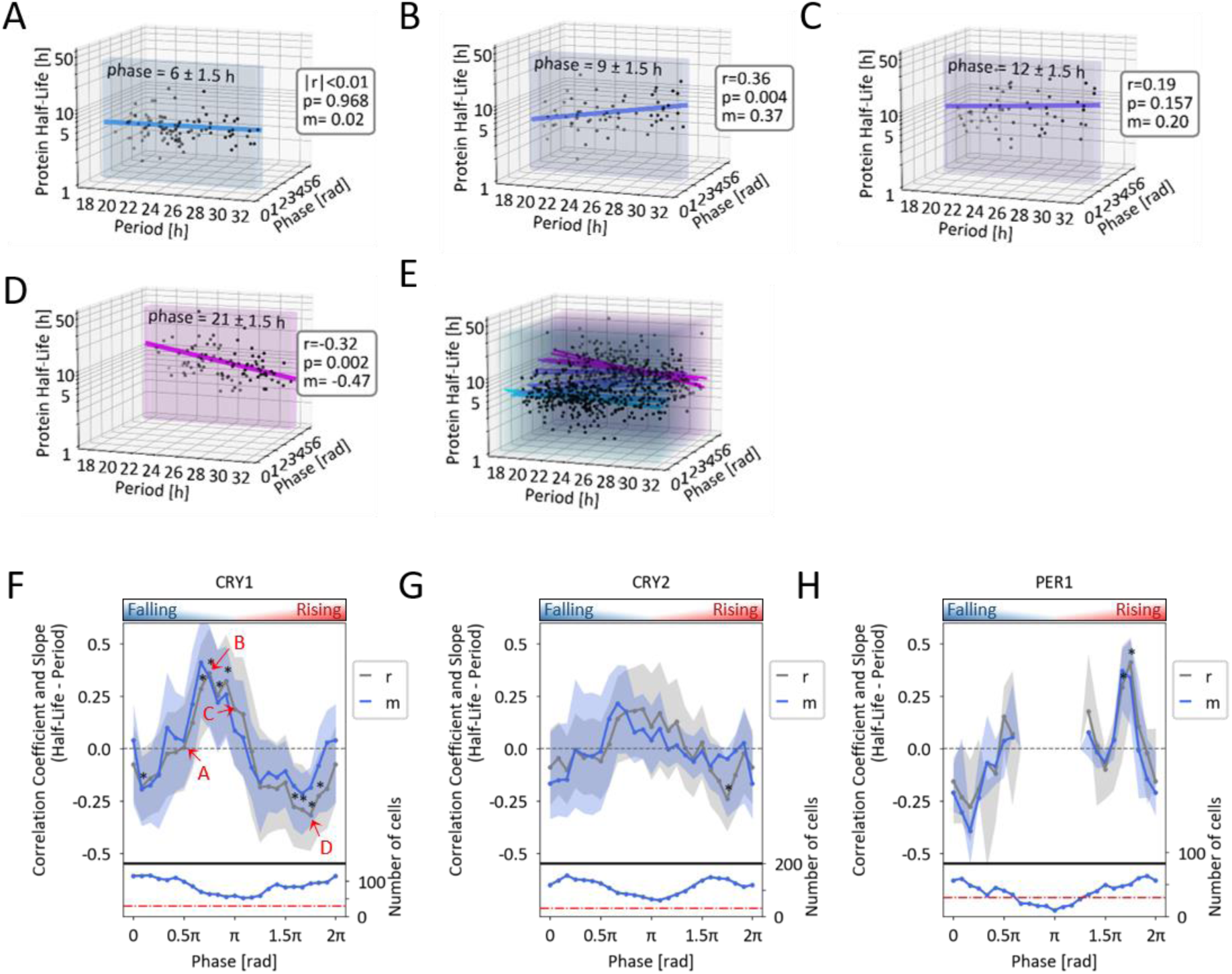
Circadian period depends on repressor protein stability in a phase-dependent manner. (A-E) 3D plots of circadian period, CRY1 half-life and circadian phase at half-life measurement. Data from indicated phase windows. (A-D) or from all phases (E) are shown. Colored lines represent linear regression (m: slope, r: Spearman correlation coefficient). (F-H) Spearman correlation coefficient and slope (m) of linear regression (median and 95% CI) for each 3-hour phase window, and number of cells for each correlation. Red dashed line: n=30 cells (minimum for correlation analysis).

In summary, CRY stability varies throughout the day, with the shortest average half-life occurring a few hours after the peak of expression and the longest half-lives occurring shortly after the trough of expression (**Fig. 3C**). However, the circadian periods of the cells seem to be rather independent of the absolute half-life of CRY1 at these extremes (**Fig. 4F-G**). In contrast, a rather short CRY1 half-life during the late rising phase (**Fig. 4G**) or a rather long CRY1 half-life during the late falling phase (**Fig. 4E**) is more often observed in cells with an above-average period and vice versa. However, even if only the stabilities assessed during the same phase are compared, cells with very different repressor half-lives can have a similar period length (**Fig. 4A-D**).

### Circadian period is compensated for protein turnover rates

One implication of the phase-dependent correlation between CRY half-life and period is that the stability of circadian repressor proteins may affect the length of the circadian cycle differentially. Intuitively, overall low protein stability could (i) prolong the time to reach a threshold of repression during the rising phase, but (ii) also shorten the time to release repression due to the accelerated disappearance of repressor proteins (**Fig. 5A**).^42^ If these effects were to cancel each other out, the period would remain stable, i.e., it would compensate for cellular fluctuations affecting protein turnover. However, intuition can easily be misled by the complexity and non-linearities present in the TTFL. Therefore, we developed an adapted mathematical model of the TTFL based on a single prototypical CRY1 repressor (**Fig. 5B**). Using linear kinetics for production terms, Michaelis-Menten kinetics for degradation terms and Hill functions for transcriptional repression, our model has four variables describing different types of the CRY1 repressor (**Supplementary Tab. S4** and Methods).

**Figure 5:**
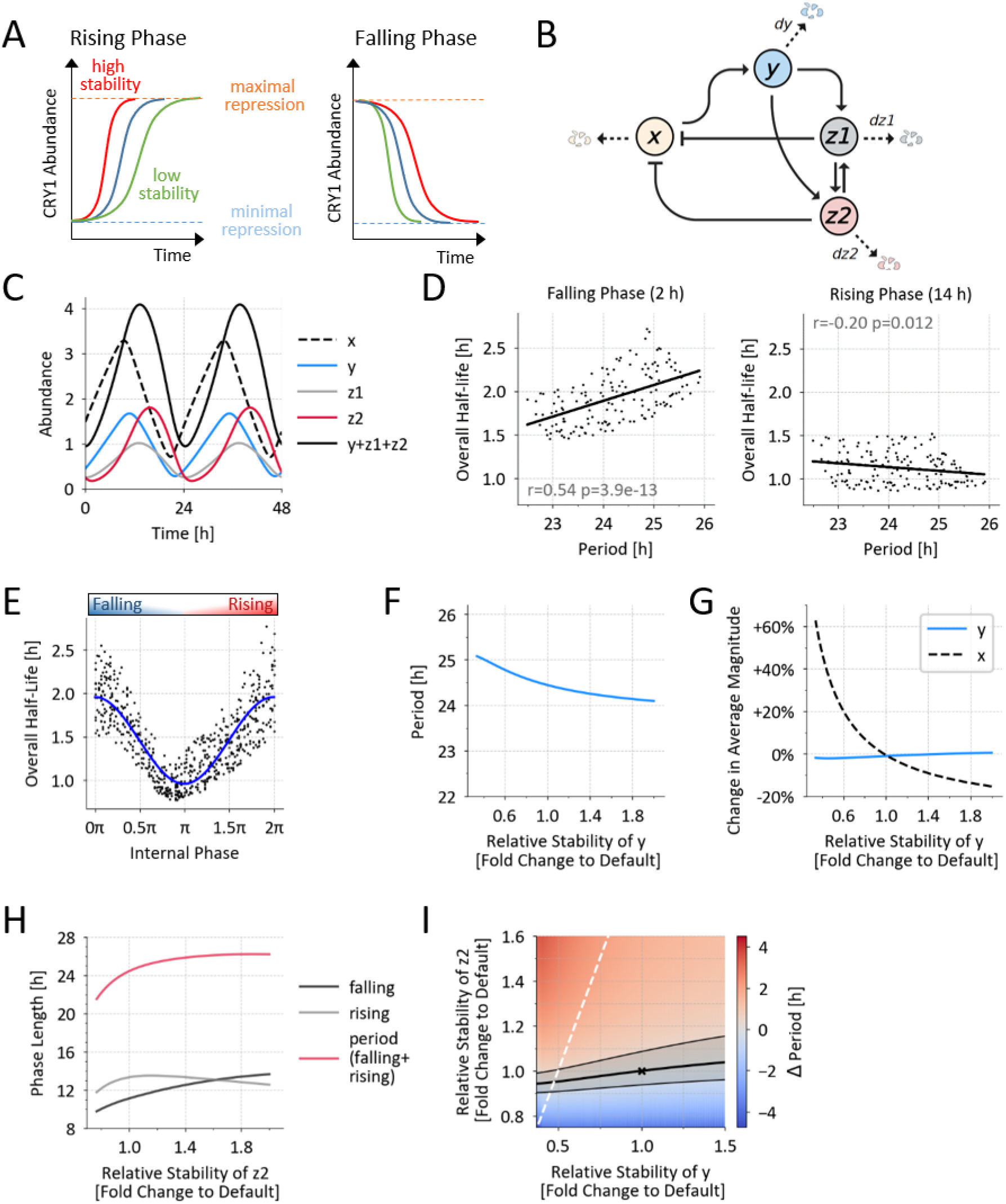
Period is compensated for protein turnover rates: a mathematical model. (A) Simple schematic of how CRY1 stability differentially affects period length. (B) Architecture of the mathematical model (see text for details). x: CRY1 mRNA, y: ‘early’, non-repressive CRY1, z1: CRY1 in high molecular weight complexes, z2: ‘late’ monomeric, repressive CRY1. (C) Oscillation in the absolute abundance of the state variables for the default parameters. (D) Regression analysis of the period and total ‘pool half-life’ (i.e., all CRY1 species y, z1, and z2) of a simulated population of single cells with stochastically varying turnover rates of early and late CRY1 for two indicated phases (n = 155). (E) Pool half-lives of the simulated population described in (D). For each cell, decay is simulated at 5 random time points. (F,G) Effect of changes in early CRY1 (y) stability on period length (F) and expression level of early CRY1 protein (y) and CRY1 mRNA (x) (G). Stability = 1/dy, dz1 and dz2 are constant. (H) Effect of changes in late CRY1 (z2) stability on length of rising phase, falling phase, and total period (rising + falling). Stability = 1/dz2, dy is constant. (I) Heatmap of period changes for different combination of y and z2 stabilities (1/y and 1/dz2, normalized to default values). Grey area shows period length of ±0.5 h from that for default parameters. White line shows where the degradation rate of y is equal to that of z2.

Transcription of this repressor leads to accumulation of mRNA (*x*) and translation into an early non-repressive CRY1 protein (*y*), which is degraded at a basal rate (*dy*). Post-translational modifications (e.g. phosphorylation) allow the repressor to inhibit its own transcription, but also make it more susceptible to degradation. This mature CRY1 (*z1* and *z2*) can either inhibit E-Box-mediated transcription in a complex (e.g. with PERs, *z1*), where it is largely shielded from FBXL3-mediated ubiquitination/degradation, or as monomeric CRY1 (*z2*), in which case it is degraded at a higher rate (degradation rate *dz2* > *dy*, *dz1*). To evaluate the total half-life of all species in this model, analogous to the experimentally measured half-life (see **Figs. 2-3**, hereafter referred to as ‘pool half-life’), translation is set to 0 and the decay curve is analyzed for 15 hours.

In this model, the mRNA and all three repressor types, as well as the total amount of protein, oscillate in a self-sustained manner (**Fig. 5C**), with the contribution of each CRY1 species to the total amount of CRY1 changing over the course of the day. While the early, non-repressive CRY1 (*y*) is the most abundant species during the accumulation phase, the late, monomeric repressive CRY1 (*z2*) dominates during the falling, repressive phase. A simulated population of single cells with stochastically varying turnover rates of early and late CRY1 (**Supplementary Fig. S7A**) shows the experimentally observed negative and positive correlation between total CRY1 half-life and period during the rising and falling phases, respectively (**Fig. 5D**). Moreover, the average of pool half-life was time-of-day dependent, as observed in our experiments (**Fig. 5E**). Thus, the model reproduced key features of our experimental data, which motivated us to take a closer look at the underlying principles.

We first investigated why the overall half-life of the CRY1 pool is time-dependent. Our model suggests two possible mechanisms, which are not mutually exclusive. First, the composition of the total CRY1 pool changes with time (from *y* to *z2*), which leads to changes in the half-life of this pool because the different CRY1 species have different degradation kinetics. The high amount of stable early CRY1 during the rising phase makes the pool more stable, whereas during the falling phase an unstable monomeric CRY1 dominates the pool composition (**Supplementary Fig. S7B**). Second, due to rate limitation of degradation processes, half-lives are longer when CRY1 abundance is high and shorter when abundance is low, resulting in a minimum pool half-life near the trough of expression (**Supplementary Fig. S7C**).

Next, we examined how the stability of the two monomeric CRY1 species affects period. We observed different effects on period depending on whether the stability of early *y* or late *z2* was changed. Increasing the stability of early CRY1 (*y*) shortens the period (**Fig. 5F**), but to a much lesser extent than we expected (**Fig. 5A**), likely due to a compensatory mechanism by reducing its production. In short, even when y stability is increased, y levels remain relatively stable because increased feedback repression leads to less mRNA (*x*) and thus less production of *y* (**Fig. 5G**), partially decoupling the dynamics of CRY1 accumulation from its stability (**Supplementary Fig. S7D**). In contrast, increasing the stability of late monomeric, repressive CRY1 (z2) lengthens the period with a saturation of the effect at very high stabilities (**Fig. 5H**). This behavior can be explained by the effect of late CRY1 stabilization on both the onset and duration of repression: While *z2* stabilization prolongs the repression, leading to a longer falling phase, it also shortens the rising phase, probably by accelerating its own accumulation, so that the threshold for repression is reached more quickly (**Fig. 5H**). These two processes have opposite effects on the period, and for stable *z2*, the shortening of the rising phase offsets the lengthening of the falling phase.

Thus, the model suggests three different mechanisms by which period may be compensated for variations in repressor stability: First, adjusted production may compensate for changes in turnover rates (e.g., of *y*). Second, changes in the stability of a particular CRY1 species (e.g., z2) may affect the length of the rising and falling phases differentially. Third, changes in the stability of different subspecies (*y* and *z2*) may have opposite effects on period length. As a consequence, different combinations of (basal and FBXL3-dependent) turnover rates may result in similar period lengths (**Fig. 5I**), and cells with the same period may have different half-lives of the CRY1 pool even when assayed at the same circadian phase (**Fig. 5D**). Thus, our model conceptualizes how a broad distribution of repressor stabilities can lead to similar circadian periods.

### FBXL3 prolongs the falling phase of CRY protein levels

One prediction of our model is that the period lengthening caused by FBXL3 knockdown, i.e., the stabilization of late CRY1, would be mainly due to an increase in the length of the falling phase of CRY1 (**Fig. 5H**). To test this, we compared the average peak shapes of thousands of normalized, peak-centered CRY1 time series recorded in the presence or absence of FBXL3 (**Fig. 6A**). Indeed, while the shape of the rising phase was little affected by FBXL3 knockdown, the average slope of the falling phase was reduced, resulting in a prolongation of the falling phase. We analyzed the distance between peak and trough at the single cell level without normalization (**Supplementary Fig. S8A**) and found that while the rising and falling phases were of similar length in wild-type cells (mean: 11.7 h vs. 12.3 h, **Fig. 6B**), FBXL3 knockdown significantly prolonged the falling phase, resulting in an asymmetric peak shape (11.9 h vs. 13.2 h, **Fig. 6B**). This was similar for CRY2 (**Supplementary Fig. S8B,E**), indicating that FBXL3 depletion indeed lengthens the period by prolonging the repressive phase, consistent with FBXL3 knockdown increasing CRY half-life mainly at times when CRY levels are decreasing (**Fig. 3F**). For PER1 and PER2, FBXL3 knockdown primarily prolonged the rising phase (**Supplementary Fig. 8C,D,F,G**). Since FBXL3 is not known to directly affect PER stability, this effect is likely caused indirectly by altered transcriptional dynamics within the TTFL.

**Figure 6:**
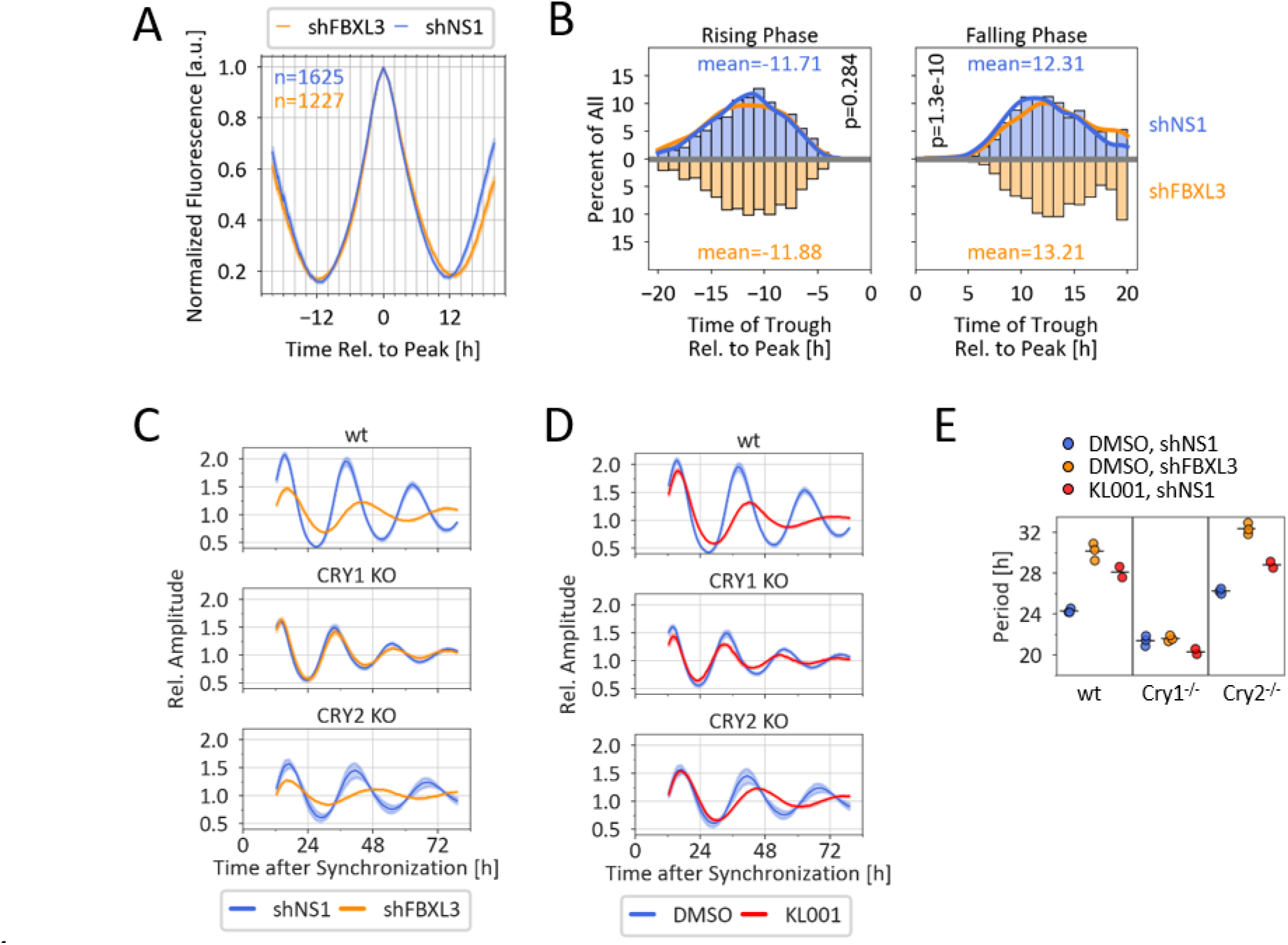
The effects of FBXL3 on circadian period is CRY1 dependent. (A) Average Peak shape of CRY1 expression after normalization (see Methods). (B) Histogram of through-to-peak durations in CRY1 time series. Lines represent kernel density estimation. p-value: Mann-Whitney-U test, n as in (A). (C,D) Detrended bioluminescence time series of wt, Cry1^-/-^ and Cry2^-/-^ *Bmal1*:Luc reporter cells after dexamethasone synchronization either transduced with non-silencing or FBXL3-targeting shRNA (C) and treated with DMSO (solvent control, (C) and (D)) or 1 µM KL001(D). Mean ±SD of 5-8 replicates representative of 3 (C) and 2 (D) experiments. (E) Periods from the recordings shown exemplarily in (C) and (D), n=2 (KL001+shNS1) and 3 (DMSO+shNS1, DMSO+shFBXL3) independent experiments, respectively.

### FBXL3 effect on period is dependent on CRY1

While depletion of FBXL3 increases the half-life of both CRY1 and CRY2, it had a greater effect on CRY1, as indicated by a greater increase in CRY1 protein half-life upon knockdown and greater loss of rhythmicity of CRY1 stability. We therefore asked whether the associated long period phenotype depends on both CRY proteins. To test this, we depleted FBXL3 in CRY1 or CRY2 knockout reporter cells^43^ and found that the period of CRY2 knockout cells and wild-type cells was lengthened by several hours after FBXL3 knockdown, but to our surprise, the period of CRY1 knockout cells was not affected (**Fig. 6C,E, Supplementary Fig. 8H**). Similar results were obtained when FBXL3 binding to CRY proteins was pharmacologically inhibited by KL001 (**Fig. 6D,E, Supplementary Fig. 8I**) suggesting that FBXL3 destabilizes CRY1 and CRY2, but its effect on circadian period is modulated primarily by its action on CRY1.

## Discussion

Circadian clocks are characterized by both robustness, e.g. the period is temperature-compensated, and plasticity, i.e. they respond to zeitgebers. While several mechanisms have been proposed for temperature compensation^44–46^, less is known about how the clock is stabilized against changes in energy supply that affect the metabolic state and thus global reaction rates such as macromolecule assembly and turnover.^47–49^

In this study, we exploit the natural heterogeneity of single cell clocks to discover fundamental principles for metabolic compensation of the circadian period without the need for genetic manipulation. By generating novel fluorescent knock-in cells targeting all major circadian repressors and simultaneously monitoring thousands of individual cells for circadian dynamics and decay characteristics, we have uncovered three key insights: (i) the length of the circadian period correlates with the stability of repressor proteins, but, contrasting the prevailing model, in a complex, phase-dependent manner; (ii) the circadian period is compensated for fluctuations in the turnover rates of circadian repressor proteins; (iii) the repressor protein stabilities are not constant, but circadian phase-dependent, and for CRY proteins this is mediated by FBXL3.

At first, we were surprised that in the naturally variable cell population, there appeared to be no dependence of circadian period on repressor protein stability, in contrast to what has often been observed.^18,23,25^ However, when analyzed separately for different phases, a complex picture emerged. First, both period length and repressor stability covered a surprisingly wide range in individual cells. Second, repressor stability and circadian period were correlated in a phase-dependent manner. Our mathematical model suggests that this is because the turnover rates of CRY subspecies are not only different, but also correlate in opposite ways with circadian period length. Depending on which species is dominant, the resulting pool half-life changes, providing an explanation for the inverse correlations: When - and only when - the stability of one species dominates the pool half-life, its true influence on period is revealed.

Thus, these opposite correlations are likely caused by the differential stabilities of the CRY subspecies and the phase-dependent differential composition of the CRY pool providing one explanation for the compensation of the period against fluctuating degradation rates. Our model suggests two additional, not mutually exclusive, mechanisms: Adapted production may counteract changes in turnover, thereby stabilizing protein levels and net flux. For example, greater CRY stability leads to greater transcriptional repression and thus less CRY production. Indeed, we and others have observed that despite increased stability, average CRY1 protein levels remain constant after FBXL3 ablation, whereas mRNA levels decrease.^19,20,50^ Finally, even changing the stability of a single CRY species can have both lengthening and shortening effects on the period, as seen for late CRY1. Thus, we propose that the circadian period is to some extent insensitive to changes in cellular protein turnover rates due to several counteracting effects.

However, mutations that affect only the turnover rate of late CRY and not the basal degradation may well affect the average period. Upon deletion of FBXL3, late CRY1 is likely to be degraded only at the basal rate (white diagonal line in **Fig. 5I**), resulting in a long period. The model predicts that even under this condition, the period is compensated for the different basal degradation rates, consistent with the lack of correlation between period and half-life observed for FBXL3 knockdown (**Supplementary Fig. S5D**). Therefore, we hypothesize that CRY mutations that prolong both CRY half-life and circadian period mainly reduce FBXL3- dependent but not basal CRY degradation.

Interestingly, we also observed that the protein half-lives of CRY1, CRY2 and PER1 are not constant, but decrease during the repression phase, paralleling previous findings on the rhythmic stability of PER2.^41^ Circadian rhythms in protein stability have long been postulated to contribute to rhythmic protein abundance. In addition to differences in translation efficiency ^51–54^, rhythmic degradation can explain rhythmic protein despite constant mRNA levels, as well as large delays between transcript and protein expression.^35,40,55–57^ Direct experimental evidence for rhythmic degradation of individual proteins is limited^41,45,58^, but our findings suggest widespread degradation rhythms^59^ consistent with circadian rhythms in autophagy and protein ubiquitination.^60,61^ For CRY1 - and to a lesser extent for CRY2 - the oscillation in protein stability depends on FBXL3, but the molecular basis is unclear. FBXL3 expression rhythms peak at times of lowest CRY1 abundance^22^ and thus cannot explain the high CRY1 stability we observed at this phase (**Fig. 3C**). Whether FBXL3 activity is regulated in a circadian manner is unknown. In addition, targeting of the substrate for ubiquitination, e.g. by post-translational modifications (PTMs) such as phosphorylation might be rhythmic. Indeed, all circadian repressors show rhythmic phosphorylation patterns suggesting that the pool of cellular repressors is not homogeneous, but consists of differentially modified subspecies^17,62^. Thus, circadian changes in degradation rates may result from changes in the composition of a pool of species with different stabilities, leading to a change in their average stability. Target accessibility may also affect degradation rates. For example, both CRY1 and CRY2 can bind to PER2 with high affinity, and the CRY-PER2 interface overlaps with the FBXL3 binding site.^9,11,63^ Thus, binding of PERs protects CRYs from degradation.^64^ A lower stability of CRYs during the falling phase could therefore be due to the absence of PERs, since CRYs are expressed later than PERs and PER concentrations at the trough are very low (**Fig. 1D**).

In CRY1-deficient U-2 OS cells, we see virtually no effect of FBXL3 knockdown or inhibition on the circadian period, in contrast to period lengthening in fibroblasts from *Cry1* knockout mice^23,65^. This suggests that in mice the long period phenotype in the absence of FBXL3 is not entirely dependent on CRY1. It is possible that this discrepancy is due to differences between mouse and human: while in mice the E3 ligase FBXL21 plays an additional competing role^66,67^, the human FBXL21 locus is a pseudogene containing a premature stop codon (NR_152421.1).^68^

## Acknowledgement

We thank all current and former members of the Kramer laboratory for technical and intellectual support. We acknowledge the support of the FCCF at the German Rheumatism Research Center. We also thank the Advanced Medical Bioimaging Core Facility (AMBIO) of the Charité for assistance with the acquisition of imaging data. We thank Christoph Harms, Steven Brown and Michela Di Virgilio for providing materials and Bharath Ananthasubramaniam and Gal Manella for helpful discussions. Adrián E. Granada’s research was supported by the German Federal Ministry of Education and Research (BMBF) through the Junior Network in Systems Medicine, under the auspices of the e:Med program (grant 01ZX1917C). This work was funded by the German Research Foundation (DFG) - project number 278001972 - TRR 186.

## Author Contribution

Conceptualization, C.H.G. and A.K.; Methodology, C.H.G., M.O. and A.K.; Software: C.H.G., M.O., A.Z., N.G.; Investigation, C.H.G., M.O., A.W., R.R., J.W., and E.H.; Writing – Original Draft, C.H.G., M.O., and A.K.; Writing – Review and Editing: C.H.G., M.O., and A.K.; Visualization: C.H.G. and M.O., Review & Editing, A.K., H.H., H.E., A.G.; Funding Acquisition, A.K., H.H.; Supervision: A.K., H.H., A.G, H.E.

## Declaration of Interest

The authors declare no competing interests.

## Methods

### Cells lines

U-2 OS (RRID:CVCL_0042, human, female, ATCC HTB-96) cells were cultured in DMEM supplemented with 10% FBS(Life, lot 2453915), 25 mM HEPES and penicillin/streptomycin at 37°C and 5% CO_2_. CRISPR KI cell lines expressing CRY1-mScarlet, CRY1-mClover and PER2- mScarlet have been described previously^4^. Cells were tested for the absence of mycoplasma using Lonza’s MycoAlert kit. For long-term imaging, cells were cultured in FluroBrite medium (GIBCO) supplemented with 2% FBS, 1x GlutaMax, 25 mM HEPES and penicillin/streptomycin from 9 days prior to imaging.

To allow automated detection of nuclei, all clones were transduced with a histone-2B-iRFP720 fusion protein, which results in nuclear expression of the infrared protein miRFP720, and cells were sorted for high expression by FACS.

### Plasmids

The generation of the original donor vectors (pDB) has been described in detail^69^ The original donor vector was modified as follows: The His/Flag tag was replaced by a 3xFLAG tag, and GSG linker sequences were introduced between protein and fluorophore and between fluorophore and 3xFLAG tag (designated pDB2). Sequences homologous to regions surrounding the stop codon of PER1 and CRY2 were synthesized by Twist bioscience and were inserted into pDB2 by restriction enzyme cloning. The pCAG-i53bp expression plasmid was a gift from Ralf Kuhn and was modified from Addgene (RRID:Addgene_74939). The SV40-NLS-CRE recombinase was a gift from Christoph Harms and was subcloned into the pLenti6 backbone. The pLenti-H2B- iRFP720 was obtained from Addgene (RRID:Addgene_128961).

Single guide RNAs (**Supplementary Tab. S3**) were designed to cut just after the STOP codon using CRISPOR^70^, and corresponding DNA oligos were ligated into pCRISPR-Lenti-v2 (RRID:Addgene_52961). To test the efficiency of the guides, cells were transduced with lentiviruses harboring the Cas9/sgRNA expression plasmid, gDNA from puromycin resistant cells was isolated, and the corresponding region was amplified by PCR and sequenced. Efficiency was assessed using the TIDE ^71^ assay.

pGIPZ clones expressing shRNA targeting FBXL3 (V2LHS_254986), CRY1 (V2LHS_172866), CRY2 (V2LHS_67009), PER1 (V2LHS_7714) or a non-silencing control (NS1), (**Supplementary Tab. S3**) were purchased from Open Biosystems (GE Healthcare) and the tGFP was mutated to abolish fluorescence. The 0.9-kb *Bmal1* promoter-driven luciferase reporter construct has been described.^42^ The sgRNA-Cas9 plasmids and donor vectors generated for this manuscript are available at Addgene (#189980-189988) along with their sequences.

### Transfection

For knock-in experiments, 10^6^ cells were harvested by trypsinization and transfected with 2 μg each of i53bp, donor vector, and pCRSIPR-Lenti-V2 by electroporation using the NEON system (Thermo Fisher, buffer N, 4 pulses, 10 ms, 1230 V). After electroporation, cells were seeded in antibiotic-free DMEM and cultured for 24 hours before selection. Transient transfections of CRE recombinase were performed using 1 μL Lipofectamine 2000 and 200 ng CRE expression plasmid in a 48-well plate format.

### Virus production and transduction of cells using lentivirus

HEK293-T cells were transiently transfected in a T75 flask with 8.6 μg lentiviral expression plasmid, 6 μg psPAX2, and 3.6 μg pMD2G (gift from Trono lab, RRID:Addgene_12259 and RRID:Addgene_12260) packaging plasmid using the CalPhos Mammalian Transfection Kit (Takara). The next day, culture medium was replaced with 12.5 mL of complete culture medium, and lentiviral supernatant was collected after 24 and 48 h. The combined supernatant was passed through a 0.45-μm filter (Filtropur S 0.45) and either used directly or stored in aliquots at −80°C. For transduction, cells were seeded into lentivirus-containing supernatant supplemented with 8 μg/mL protamine sulfate. The next day, lentivirus-containing supernatant was aspirated and cells were cultured in complete culture medium for another 24 hours before antibiotic selection of transduced cells.

### Antibiotic selection

To select for transfected or transduced cells, cells were grown subconfluently in medium containing blasticidin (10 μg/ml) for >3 days or in medium containing puromycin (10 μg/ml) for >1 day until non-transfected control cells died.

### FACS sorting

Cells were sorted on a FACS AriaII (BD). For staining of surface hCD4 for negative selection, 2 × 10^6^ cells were trypsinized, washed with 0.5% BSA/ PBS and incubated with 200 μL of a 1:50 dilution of hCD4-BV711 (OKT4, Bio-Legend, UK) for 30 minutes. The cells were washed twice with BSA/PBS. Excitation: 405 nm. Emission filter: 525LP-525/50 (CFP), 685LP-710/50 (BV711).

### Nucleic acid isolation and PCR

Genomic DNA was extracted using Direct PCR Lysis Reagent Cell (VWR). RNA was extracted using the AMBION PureLink RNA Mini Kit (Thermo Fisher) according to the manufacturer’s instructions, including an on-column DNase digest. RNA was reverse transcribed using a primer that anneals to the 3xFLAG sequence in a two-step protocol. PCR amplification was performed with Phusion polymerase (New England Biolabs), and products were analyzed by agarose gel electrophoresis and detected using RedSafe/UV light. Primer sequences are listed in **Supplementary Tab. S3**.

### Bioluminescence recording of circadian oscillations

Cells transduced with a reporter plasmid in which luciferase expression is driven by a *mBmal1* promoter fragment, were seeded to confluence. To synchronize circadian rhythms, cells were treated with 1 μM dexamethasone for 20 minutes followed by two washes with warm PBS. Cells were then incubated in DMEM without phenol-red supplemented with 250 μM D-luciferin, and dishes were sealed with parafilm. Bioluminescence was recorded in a LumiCycle (Actimetrics) or TopCount (Perkin Elmer). Raw data were detrended by dividing by the 24-hour running average. Periods, phases and mean bioluminescence signal were estimated by fitting the cosine wave function using ChronoStar software.

### Fluorescence microscopy

For microscopy, cells were seeded on glass bottom #1.5H-N 96-well plates (Cellvis, USA) coated with 50 µg/ml human serum fibronectin (Merck, Germany). Imaging was performed on a Nikon Widefield Ti2 equipped with a sCMOS, PCO.edge camera and a live cell incubator. Images were acquired in Flurobrite medium (GIBCO) supplemented with 2% FBS, 1:100 PenStrep, and 1x GlutaMax at 37°C and 5% CO2. The following light sources (LEDs) and emission filters were used for the different channels: YFP (mClover3): excitation 511/16 nm, 12.3 mW, 30% intensity, emission 540/30 nm; RFP (mScarlet-I): excitation 555/28 nm, 145 mW, 12% intensity, emission 642/80 nm; iRFP: excitation 635/22 nm, 38.9 mW, 75% intensity, emission 697/60 nm. Objectives: 40x ApoFluor, NA 0.95, WD 250 μm. Illumination time for iRFP 700 ms and 2 s for all other channels. Images were acquired in a regular imaging interval of 1 h.

### Cell tracking and quality control

Cell tracking was performed automatically using Cellprofiler. Multidimensional .nd2 files were decomposed into individual tiff files using Fiji. Per channel, 100 images from buffer only wells were loaded into Cell Profiler (pipeline 1, supplemental material) and used to generate relative illumination patterns for each channel. Images from each time series were loaded into Cellprofiler (pipeline 2, modified from Manella et al.^72^, supplemental material). Within this pipeline, images were corrected for non-uniform illumination by dividing pixel by pixel by the previously generated patterns. iRFP channel was used for segmentation of nuclei in each image and subsequent tracking of nuclei throughout the time series. The background of the illumination-corrected RFP and YFP images was determined by the median fluorescence intensity of all unsegmented pixels (i.e. not identified as nuclei). Finally, the mean fluorescence intensity for each tracked nucleus for each time point was extracted from the RFP and YFP channel. After cell division, tracking continued with one daughter cell, while the other daughter cell was considered a newly emerging object. Only objects tracked for at least 24 subsequent images were retained at this stage (primary objects).

For quality control, we developed a Python script (Note: script will be made publicly available on GitHub) that detects abrupt changes in nuclear size of >20% and cell division events, defined as a peak in average H2B-iRFP720 fluorescence due to chromatin condensation, followed by a decrease (>20%) in nuclear size. Subsequently, all size changes not related to cell divisions were flagged as potential tracking/segmentation errors. Time series were cropped to exclude errors and accepted if they contained ≥60 error-free consecutive images. Overall, 9-31% of primary objects passed these quality control criteria. We visually inspected a subset of accepted time series and estimated that ∼90% were correctly tracked. Fluorescence intensities at cell division and subsequent time points were linearly extrapolated from neighboring time points, because detachment of dividing cells produced fluorescence artifacts.

### Circadian parameter extraction and rhythmicity threshold

Circadian parameters were determined using metacycle2D^36^ with LS and JTK cycle analysis, a period range of 18-32 h and Fisher corrected p-values. Where appropriate, input data were truncated to begin 24 hours after synchronization or to end at the time of CHX addition. Phase at CHX addition was calculated from phase and period using a equation (E1):

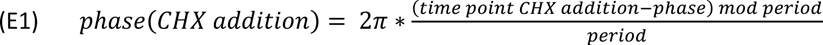

To calculate a high-confidence threshold for rhythmicity for each channel, time series of non-fluorescent cells recorded during the same experiment were analyzed in parallel, and the threshold was set to the 5^th^ percentile of the p-value from these time series. For a time series to be considered rhythmic, its p-value had to exceed this threshold. See **Supplementary Note SN1** for details.

### Determination of photobleaching

Prior to each experiment, photobleaching was measured by imaging cells from different clones 20 times within 1 hour using the same microscope settings, a time frame in which signal decay is expected to be dominated by photobleaching. Cells were tracked and monoexponential decay curves fitted to time point 3-16 of the individual cell time series using the Python package *scipy.optimze.curve_fit* and equation (E2):

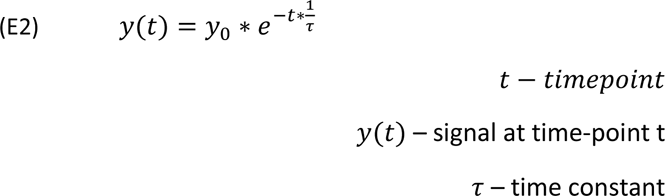

and filtered for fits with a correlation coefficient r²>0.7. The median time constant τ was then calculated for each fluorophore.

### Calculation of protein half-life

Protein half-life was calculated from time points 2-8 hours after CHX addition. For each channel, background was determined as the mean intensity of non-fluorescent nuclei. We excluded data from cells whose intensity at time point 2 h did not clearly exceed background (median + 2*SD). Median background was subtracted from all time series and time series were corrected for additive photobleaching using equation (E3):

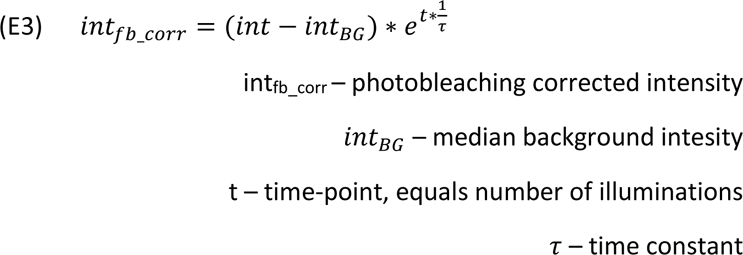

Decay parameters were calculated by fitting monoexponential decay curves (no plateau) to the photobleach-corrected time series after CHX addition as described above, and filtering for fits with a correlation coefficient r² of >0.7. Finally, the half-life was calculated using equation (E4)

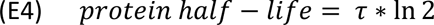

### Analysis of phase length and average peak shapes

Rhythmic fluorescence time series were smoothed by calculating the running average of 3 consecutive time points. Peak time was determined as the maximum intensity within 5 hours of the first calculated peak phase time (Metacylce2D), and trough time was determined as the minimums within 20 hours either before or after the peak time. The length of rising and falling phases was determined as the time difference between peak and trough times.

For extraction of average peak shapes, signal intensities of rising and falling phases were further normalized independently between 0 and 1. The time axes of the time-series were stretched to the median period of each genotype, and the average peak shape was calculated as mean±SEM.

### Mathematical modelling

We have developed an adapted mathematical model of the transcription-translation feedback loop (TTFL) based on a single CRY1 repressor, which is based on the classical model described by Goodwin more than 50 years ago.^73^ Our model was developed to capture the dual inhibition mechanism of CRY1: in the earlier phase of repression, CRY1 interacts with other PER and CRY proteins to form a high molecular weight complex that binds and inhibits the activator complex containing CLOCK and BMAL1^9–12^. In the later circadian repression phase, CRY1 independently represses E-box-induced transcription.^13,14,74,75^

We used linear terms to model production and import/export terms, Michaelis-Menten kinetics for degradation processes and Hill functions with an ‘AND’ funnel ^76^ for both modes of transcriptional repression. The model equations are given below as (E5-E8):

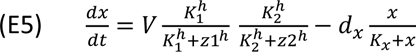

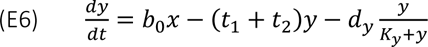

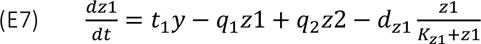

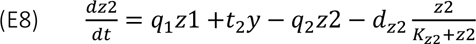

With the above assumptions (dual inhibition mechanism of CRY1, and linear, Michaelis-Menten and Hill kinetics to describe the biological processes), we systematically explored the parameter space to find sets of parameters that reproduce our experimental findings, namely (i) the phase-dependent stability of total CRY1 protein (y+z1+z2) (**Fig. 3**); (ii) a later peak phase of z2 than that of z1 (**Fig. 1L**); (iii) a positive correlation of the oscillator period with the overall stability of total CRY1 protein during the falling phase (**Fig. 4B**); and (iv) a negative correlation of the circadian period with the overall stability of total CRY1 protein (y+z1+z2) during the rising phase (**Fig. 4D**). Stability of total CRY1 (y+z1+z2) was calculated by setting the translation rate (b0) to 0 (mimicking CHX addition) and fitting a mono-exponential decay function to the decay curve of total CRY1 protein (y+z1+z2). The pool half-life was calculated from the fitted parameters.

The default values of the wild-type parameters are listed in **Supplementary Tab. S4**. These values were chosen to demonstrate the plausibility of our conceptual model and should not be considered as exact representations of the true biochemical rate constants. To simulate cell-to-cell heterogeneity, we randomly varied the turnover rates of early and late CRY1 (**Supplementary Fig. S6A**). The turnover rates of y were drawn from a uniform distribution, allowing dy to vary between 20% and 300% of its default parameter value. In the case of z2, we limited the range of variation to 80% to 120% of its default parameter value, as large changes in dz2 resulted in the loss of oscillations. We also ensured that dz2 was at least as large as the basal degradation rate dy. Numerical simulations were performed in Python using the *odeint* function from the *scipy* library to solve the ordinary differential equations.

### Blinding and randomization, data exclusion, statistical analysis

For two of the three experimental runs (run 1 and 2, **Supplementary Tab. S5**), the virus type (shRNA) was blinded to the experimenter. Prior to seeding cells into the 96-well plate, clonal identity was blinded to the experimenter, which also meant that cell seeding was randomized. Deblinding was performed during automated quality control and no data were manually excluded afterwards, with the exception of one clone in experiment 2 that did not show the expected fluorescence. During data analysis, non-rhythmic time series were excluded from all analyses requiring determination of circadian periods, phases, or amplitudes. Protein half-lives derived from poor fits (r²<0.70) or low initial intensities were ignored, and corresponding time series were excluded from all analyses requiring determination of protein half-life. Statistical analyses were performed in Python using the *scipy* library. Unless otherwise noted, all statistical tests were two-sided.

## Data and material availability

Imaging raw and metadata will be provided via the EMBL-EBI BioImage Archive (https://www.ebi.ac.uk/bioimage-archive) with accession number ###. The Cellprofiler pipelines used for analysis are deposited on GitHub along with supporting and output files. All own Python scripts can be found at GitHub (###). A data table of all successfully tracked cells including raw data and derived values is included in the supplement (**Supplementary Tab. S5**). All relevant plasmids are deposited at Addgene (#179441, 179453, 189980-189988). All cell lines generated in this study are available from the lead contact with a completed Material Transfer Agreement.

## Notes

### Competing Interest Statement

The authors have declared no competing interest.

