## Supplementary Information for "Circadian period is compensated for repressor protein turnover rates in single cells"

Supplementary Note S1

Supplementary Tables

Supplementary Figures (including Supplementary Figure Legends)

Supplementary Video Legends

#### Supplementary Note SN1:

We used Metacylce2D to identify rhythmic time-series with a period range of 18-32 hours. Metacycle2D identified circadian rhythms in over 90% of the time-series with a p-value < 0.05 percent. To exclude direct effects of dexamethasone, the analysis was restricted to time points starting 24 hours after synchronization. We wondered whether fluctuations in auto-fluorescence or experimental conditions may in part be responsible for the high percentage of rhythmic time-series. To address this issue, we also searched for rhythms in time-series of background-subtracted nuclear auto-fluorescence, obtained from cells that do not express a fluorophore in the respective channel. Strikingly, more than 20% of these time-series were determined rhythmic by Metacycle2D with a p-value < 0.05. To focus our analysis only on time-series in which the reporter signal oscillates well beyond these fluctuations, we calculated the 5$^{th}$ percentile of all p-values from the auto fluorescence time-series of a dataset (~$2*10^{-6}$ and ~$2*10^{-8}$ for the dataset related to Fig.1 and Fig. 2-6, resp.). We used this value as a cutoff for rhythmicity, and demanded that the Metacycle2D p-value of a fluorescence time-series had to surpass this threshold to be considered as rhythmic. By increasing stringency, we also increased confidence in the calculated circadian parameters for further analysis, at the expense of potentially excluding a part of rhythmic time- series, especially those with lower absolute amplitudes. We furthermore excluded times-series which were considered as rhythmic, but whose calculated period matched exactly the inputted limits (18-32h), as these most probably contain oscillations longer or shorter than these limits.

### Supplementary Tables

**Supplementary Table S1**: Clones used for live cell imaging.

| Cell clone | mClover3 | mScarlet-I | Parental clone | First described |
|---|---|---|---|---|
| **CRY1-mClo #1** | CRY1, biallelic | --- | Wild-type | Gabriel et al., 2021[4] |
| **CRY1-mClo/PER2-mSca #6** | CRY1, monoallelic | PER2, monoallelic | PER2-mSca #2 | Gabriel et al., 2021[4] |
| **PER2-mSca #1** | --- | PER2, monoallelic, large deletion second allele | Wild-type | Gabriel et al., 2021[4] |
| **CRY1-mSca #1** | --- | CRY1, monoallelic | Wild-type | Gabriel et al., 2021 [4] |
| **CRY2-mSca #1** | --- | CRY2, biallelic * | Wild-type | This report |
| **CRY2-mSca #2** | --- | CRY2, biallelic * | Wild-type | This report |
| **PER1-mSca #1** | --- | PER1, monoallelic* | Wild-type | This report |
| **PER1-mSca #2** | --- | PER1, monoallelic* | Wild-type | This report |
| **CRY1/CRY2 DKI #1** | CRY2, monoallelic* | CRY1, biallelic | CRY1-mCl3 #1 | This report |
| **CRY1/CRY2 DKI #2** | CRY2, monoallelic* | CRY1, biallelic | CRY1-mCl3 #1 | This report |
| **CRY1/CRY2 DKI #3** | CRY2, biallelic * | CRY1, monoallelic* | CRY2-mSca #2 | This report |
| **CRY1/CRY2 DKI #4** | CRY2, biallelic * | CRY1, monoallelic* | CRY1-mCl3 #2 | This report |

*classified as monoallelic / biallelic based on presence or absence of the wild-type band in genomic PCR

**Supplementary Table S2**: Overview of nuclear fluorescence time series

| Cell clone | Virus | Protein | Tracked objects that passed QC | Rhythmic time series | Rhythmic [% of all] | Accepted decay fit | Accepted fit [% of all] | Accepted fit + rhythmic | Accepted fit + rhythmic [% of all] | Accepted fit + rhythmic [% of rhythmic] |
|---|---|---|---|---|---|---|---|---|---|---|
| **CRY1-mClo** | **shFBXL3** | **CRY1** | 541 | 134 | 24,8% | 300 | 55,5% | 71 | 13,1% | 53,0% |
| | **shNS1** | **CRY1** | 539 | 241 | 44,7% | 340 | 63,1% | 154 | 28,6% | 63,9% |
| **CRY1-mSca** | **shFBXL3** | **CRY1** | 807 | 179 | 22,2% | 628 | 77,8% | 150 | 18,6% | 83,8% |
| | **shNS1** | **CRY1** | 948 | 330 | 34,8% | 742 | 78,3% | 263 | 27,7% | 79,7% |
| **CRY1/2 DKI #1** | **shFBXL3** | **CRY1** | 383 | 93 | 24,3% | 146 | 38,1% | 33 | 8,6% | 35,5% |
| | | **CRY2** | 383 | 87 | 22,7% | 256 | 66,8% | 62 | 16,2% | 71,3% |
| | **shNS1** | **CRY1** | 263 | 74 | 28,1% | 93 | 35,4% | 36 | 13,7% | 48,6% |
| | | **CRY2** | 263 | 80 | 30,4% | 108 | 41,1% | 35 | 13,3% | 43,8% |
| **CRY1/2 DKI #2** | **shFBXL3** | **CRY1** | 570 | 160 | 28,1% | 226 | 39,6% | 60 | 10,5% | 37,5% |
| | | **CRY2** | 570 | 96 | 16,8% | 363 | 63,7% | 70 | 12,3% | 72,9% |
| | **shNS1** | **CRY1** | 757 | 200 | 26,4% | 342 | 45,2% | 95 | 12,5% | 47,5% |
| | | **CRY2** | 757 | 183 | 24,2% | 401 | 53,0% | 112 | 14,8% | 61,2% |
| **CRY1/2 DKI #3** | **shFBXL3** | **CRY1** | 504 | 111 | 22,0% | 137 | 27,2% | 28 | 5,6% | 25,2% |
| | | **CRY2** | 504 | 117 | 23,2% | 411 | 81,5% | 94 | 18,7% | 80,3% |
| | **shNS1** | **CRY1** | 544 | 180 | 33,1% | 200 | 36,8% | 69 | 12,7% | 38,3% |
| | | **CRY2** | 544 | 224 | 41,2% | 349 | 64,2% | 155 | 28,5% | 69,2% |
| **CRY1/2 DKI #4** | **shFBXL3** | **CRY1** | 490 | 99 | 20,2% | 241 | 49,2% | 54 | 11,0% | 54,5% |
| | | **CRY2** | 490 | 107 | 21,8% | 319 | 65,1% | 71 | 14,5% | 66,4% |
| | **shNS1** | **CRY1** | 212 | 61 | 28,8% | 61 | 28,8% | 20 | 9,4% | 32,8% |
| | | **CRY2** | 212 | 60 | 28,3% | 78 | 36,8% | 21 | 9,9% | 35,0% |
| **CRY1/PER2 DKI** | **shFBXL3** | **CRY1** | 295 | 66 | 22,4% | 108 | 36,6% | 30 | 10,2% | 45,5% |
| | | **PER2** | 295 | 25 | 8,5% | 1 | 0,3% | 0 | 0,0% | 0,0% |
| | **shNS1** | **CRY1** | 249 | 74 | 29,7% | 135 | 54,2% | 37 | 14,9% | 50,0% |
| | | **PER2** | 249 | 22 | 8,8% | 7 | 2,8% | 0 | 0,0% | 0,0% |
| **CRY2-mSca #1** | **shFBXL3** | **CRY2** | 467 | 164 | 35,1% | 369 | 79,0% | 130 | 27,8% | 79,3% |
| | **shNS1** | **CRY2** | 579 | 261 | 45,1% | 386 | 66,7% | 190 | 32,8% | 72,8% |
| **CRY2-mSca #2** | **shFBXL3** | **CRY2** | 1490 | 406 | 27,2% | 886 | 59,5% | 254 | 17,0% | 62,6% |
| | **shNS1** | **CRY2** | 1520 | 750 | 49,3% | 714 | 47,0% | 376 | 24,7% | 50,1% |
| **PER1-mSca #1** | **shFBXL3** | **PER1** | 383 | 181 | 47,3% | 74 | 19,3% | 44 | 11,5% | 24,3% |
| | **shNS1** | **PER1** | 527 | 316 | 60,0% | 231 | 43,8% | 129 | 24,5% | 40,8% |
| **PER1-mSca #2** | **shFBXL3** | **PER1** | 897 | 383 | 42,7% | 201 | 22,4% | 100 | 11,1% | 26,1% |
| | **shNS1** | **PER1** | 833 | 413 | 49,6% | 341 | 40,9% | 166 | 19,9% | 40,2% |
| **PER2-mSca** | **shFBXL3** | **PER2** | 1838 | 192 | 10,4% | 12 | 0,7% | 3 | 0,2% | 1,6% |
| | **shNS1** | **PER2** | 1268 | 229 | 18,1% | 5 | 0,4% | 1 | 0,1% | 0,4% |

**Supplementary Table S3:** DNA sequence information

| Name | Description / Figure | Sequence |
|---|---|---|
| **sgRNA CRY2** | | AGCAGCCTAGATTCAACCTC |
| **sgRNA PER1** | | GAAAGCCTCTCATGGACTCC |
| **sgRNA CRY1** | | GAAACGTCCTAGTCAGGAAG |
| **V2LHS_254986** | shRNA hFBXL3 | tgctg ttgac agtga gcgcg GACAT TATTC TCCAA GTATt tagtg aagcc acaga tgtaa atact tggag aataa tgtcc ttgcc tactg cctcg ga |
| **V2LHS_172866** | shRNA hCRY1 | tgctg ttgac agtga gcgcg CTGAG GCAAG CCGTT TGAAt tagtg aagcc acaga tgtaa ttcaa acggc ttgcc tcagc atgcc tactg cctcg ga |
| **V2LHS_67009** | shRNA hCRY2 | tgctg ttgac agtga gcgac GCTTG ACCTT TGAAT ATGAc tagtg aagcc acaga tgtag tcata ttcaa aggtc aagcg gtgcc tactg cctcg ga |
| **V2LHS_7714** | shRNA hPER1 | Tgctg ttgac agtga gcgag CCTTG GTGCT CCCTA ACTAt tagtg aagcc acaga tgtaa tagtt aggga gcacc aaggc ctgcc tactg cctcg ga |
| **Primer CRY1 genome fw** | Fig. S4C | actgccactgattgcctgggattgaagt |
| **Primer CRY1 genome rv** | Fig. S4C | cagctgcaacagtattcctcctg |
| **Primer CRY2 genome fw** | Fig. S1D, Fig. S4B | ggctggcagcatgagcagtg |
| **Primer CRY2 genome rv** | Fig. S1D, Fig. S4B | gcggcctggccctgaagta |
| **Primer PER1 genome fw** | Fig. S1E | ctgagcctggggaccaggtg |
| **Primer PER1 genome rv** | Fig. S1E | cctcagggcaccacggagtt |
| **Primer 3xFLAG rv** | RT-primer, Fig. S1B-C | ttgtcatcgtcatccttgtaatcg |
| **Primer mScarlet-I cDNA rv** | Fig. S1B-C | gtcttgaagtccgccaggtagc |
| **Primer CRY2 cDNA fw** | Fig. S1B | gccctggaatgccccagagt |

**Supplementary Table S4:** Default parameters of the mathematical model

| Parameter | Default Value | Interpretation |
|---|---|---|
| $\mathbf{b_0}$ | 0.35 $h^{-1}$ | CRY1 translation rate |
| $\mathbf{t_1}$ | 0.40 $h^{-1}$ | rate of macromolecular complex formation |
| $\mathbf{t_2}$ | 0.10 $h^{-1}$ | nuclear import rate of monomeric CRY1 |
| $\mathbf{q_1}$ | 0.75 $h^{-1}$ | macromolecular complex dissociation rate |
| $\mathbf{q_2}$ | 0.15 $h^{-1}$ | macromolecular complex association rate |
| $\mathbf{d_x}$ | 0.35 [conc.]/$h^{-1}$ | degradation rate of *CRY1* mRNA (x) |
| $\mathbf{d_y}$ | 0.30 [conc.]/$h^{-1}$ | degradation rate of early non-repressive CRY1 (y) |
| $\mathbf{d_{z1}}$ | 0.15 [conc.]/$h^{-1}$ | degradation rate of early repressive CRY1 complex (z1) |
| $\mathbf{d_{z2}}$ | 0.60 [conc.]/$h^{-1}$ | degradation rate of late monomeric CRY1 (z2) |
| **V** | 0.5 [conc.]/$h^{-1}$ | E-box activation rate |
| $\mathbf{K_1}$ | 1.0 [conc.] | $EC_{50}$ for transcriptional inhibition by z1 |
| $\mathbf{K_2}$ | 1.0 [conc.] | $EC_{50}$ for transcriptional inhibition by z2 |
| **h** | 8 a.u. | Hill coefficient for transcriptional repression |
| $\mathbf{K_x}$ | 0.5 [conc.] | $K_M$ constant for x degradation |
| $\mathbf{K_y}$ | 0.25 [conc.] | $K_M$ constant for y degradation |
| $\mathbf{K_{z1}}$ | 1.0 [conc.] | $K_M$ constant for z1 degradation |
| $\mathbf{K_{z2}}$ | 0.25 [conc.] | $K_M$ constant for z2 degradation |

**Supplementary Table S5:** Single cell data (separate file including data description).

### Supplementary Figures

#### Supplementary Figure S1

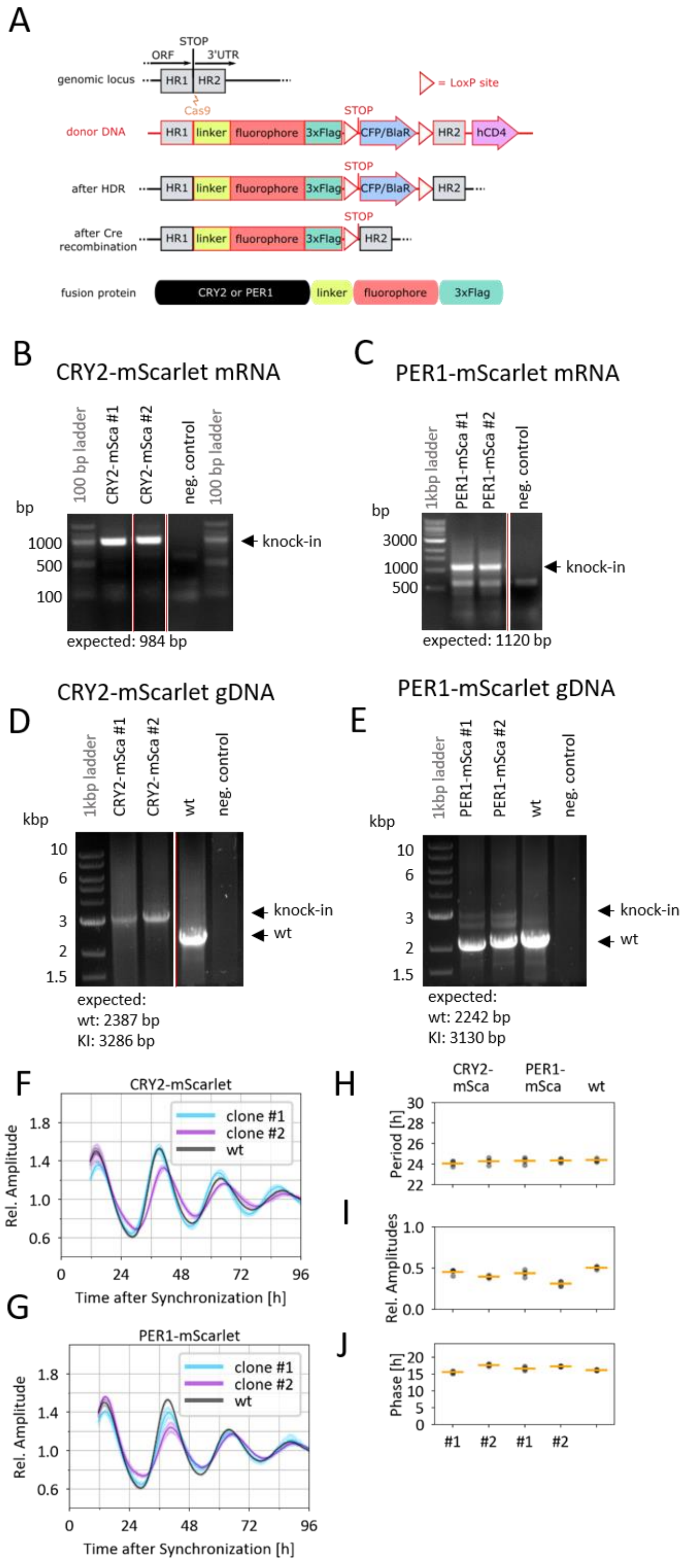

#### Supplementary Figure S1:

(A) Knock-in strategy to fluorescence-tagged proteins C-terminal to the endogenous locus. HR: homology region, HDR: homology directed repair, UTR: untranslated region.

(B-C) Chimeric mRNA was detected in knock-in clones by reverse transcription-polymerase chain reaction (RT-PCR) using a RT and a reverse primers specific for the inserted sequence and a gene-specific forward primer.

(D-E) Successful knock-in was confirmed by PCR amplification of the targeted region of the CRY2 (D) and PER1 (E) genomic loci, showing either the wild-type allele, the larger knock-in allele or both. Additional intermediate bands in (E) are most likely heteroduplexes of wild-type and knock-in products.

(F-G) Detrended bioluminescence time series of individual CRY2 (F) and PER1 (G) knock-in clones and wild-type transduced with a *Bmal1*:Luc reporter after synchronization by dexamethasone. Mean + SD of 4 measurement cohorts from 2 independent experiments, each consisting of 4-8 replicates (n=4). (H-J): Extracted periods (H), amplitudes (I) and circadian phase (J) from D and E (n=4).

### Supplementary Figure S2

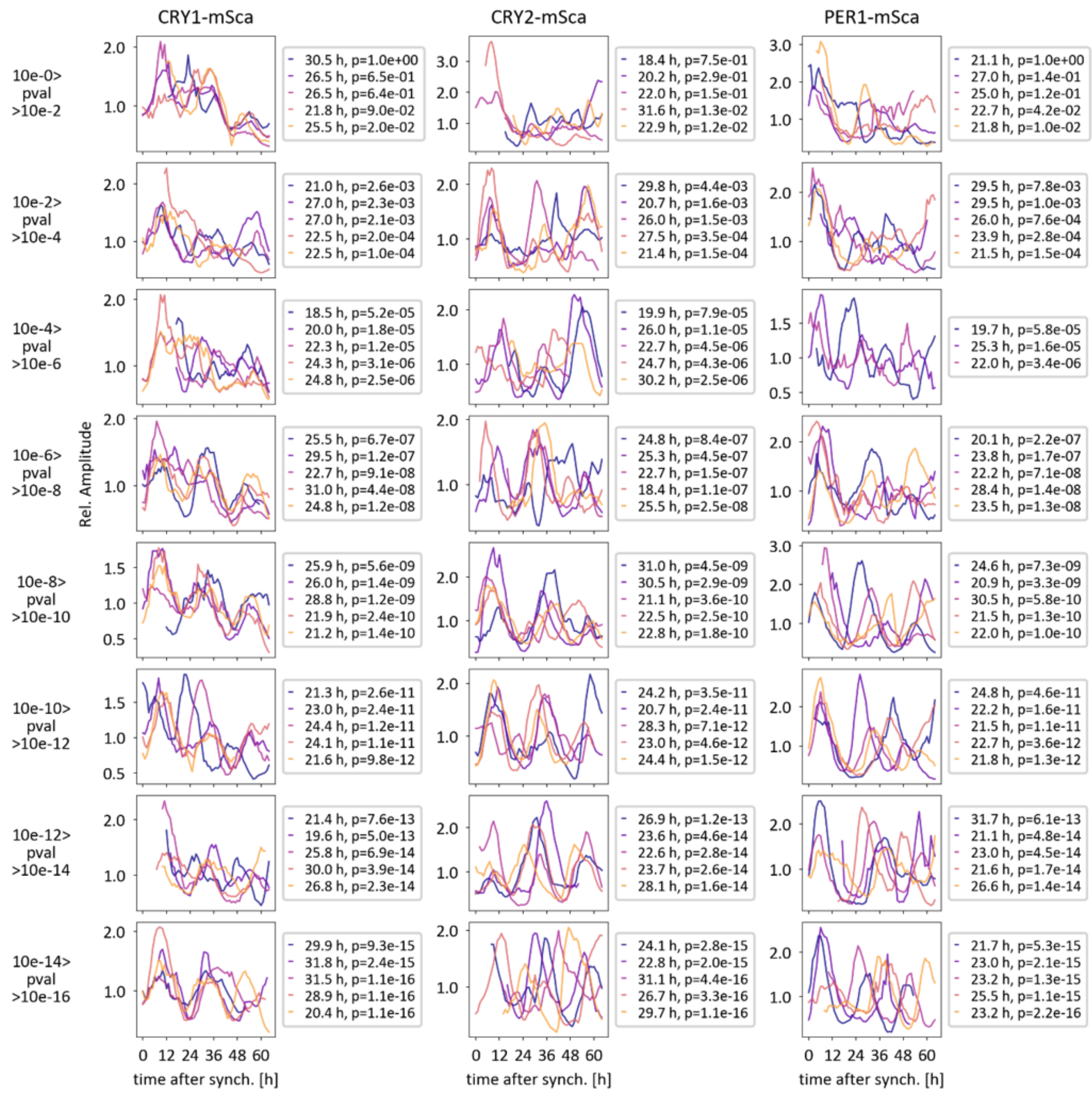


Supplementary Figure S2:

Examples of background-subtracted and normalized nuclear fluorescence time series of indicated knock-in clones from Fig. 1E. Signals grouped according to Metacycle2D-derived p-values for rhythmicity.Period length and p-value are provided in figure legends.

### Supplementary Figure S3:

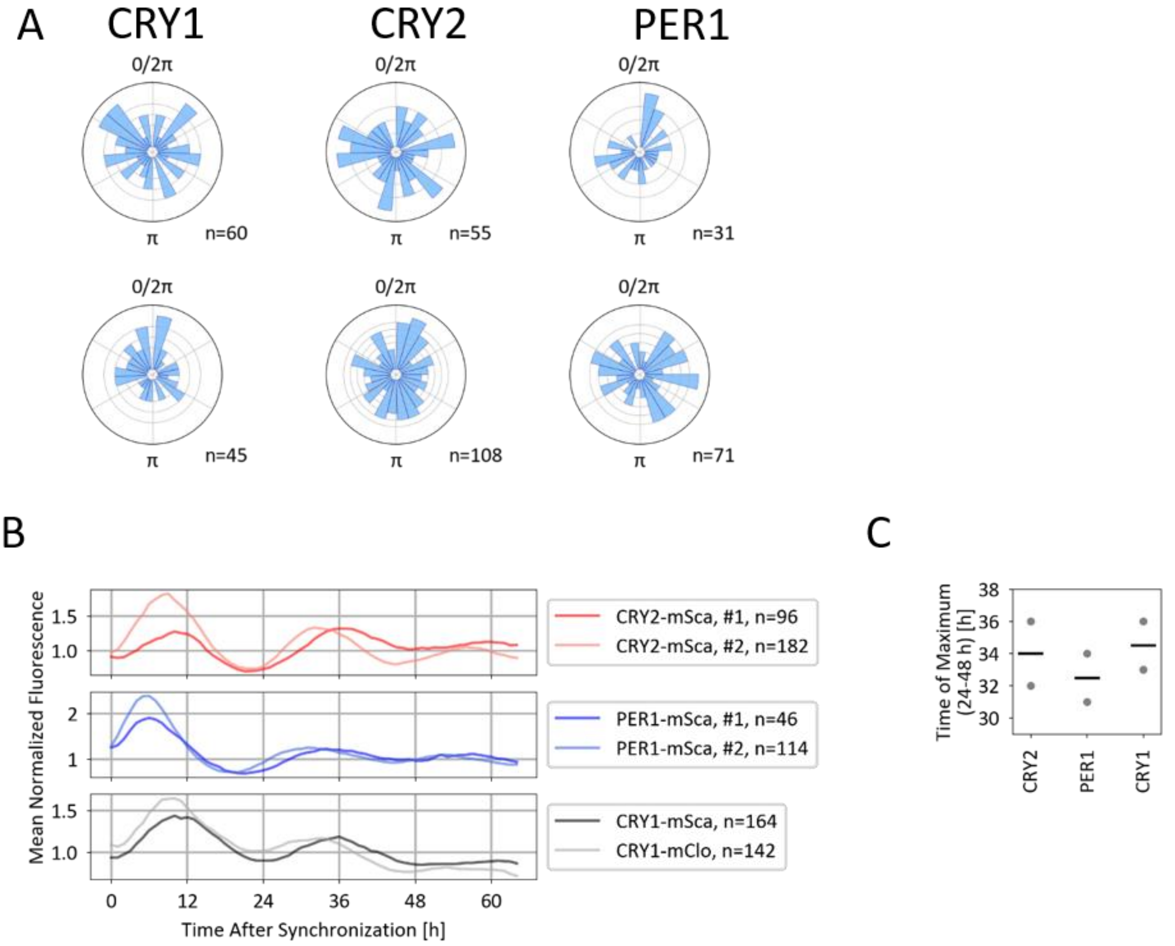


Supplementary Figure S3:

(A) Histogram of circadian phases 24-65 hours after synchronization of rhythmic time series from Fig. 1E, 1-h bins, 1 cell per ring.

(B) Population mean of normalized fluorescence time series per clone.

(C) Clonal and mean peak time of fluorescence intensity on day 2 after dexamethasone treatment (derived from B).

### Supplementary Figure S4:

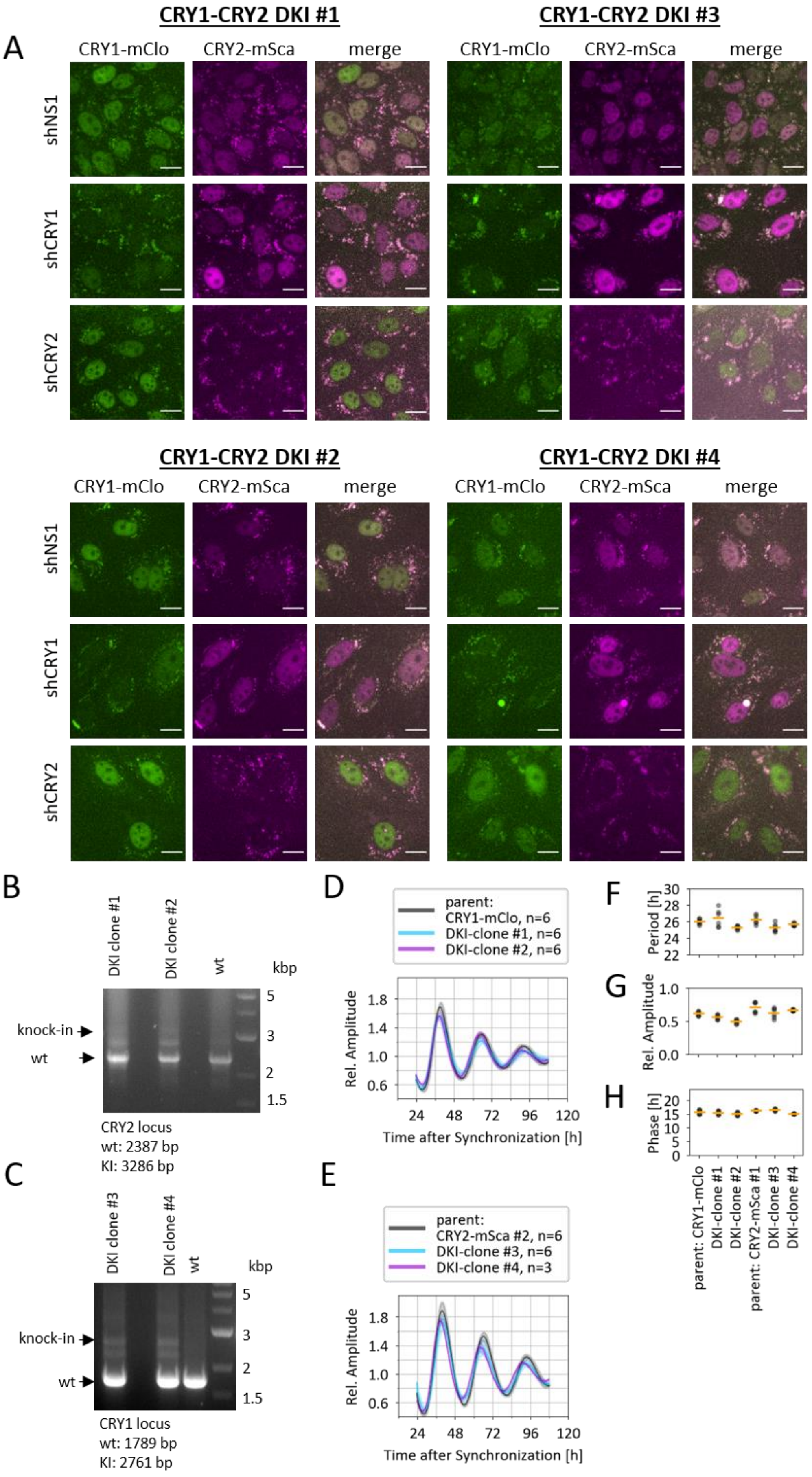

#### Supplementary Figure S4:

(A) Fluorescence images of selected CRY1/CRY2 double knock-in clones. Cells were transduced with shRNA targeting either CRY1 or CRY2, or with a non-silencing shRNA. Scale bar = 20 µm.
(B-C) Successful knock-in was confirmed by PCR amplification of the targeted region of CRY2 (B) and CRY1 (C) genomic loci, showing either wild-type allele, the larger knock-in allele or both. Additional intermediate bands are most likely heteroduplexes of wild-type and knock-in products.
(D-E) Detrended bioluminescence time series of CRY1-CRY2 double knock-in and parental single knock-in clones transduced with a *Bmal1*:Luc reporter. Mean + SD of 3-6 replicates from 1 (DKI clone 4) or 2 independent experiments (n=3-6).
(F-H) Extracted periods (F), amplitudes (G) and circadian phase (H) from (D) and (E).

### Supplementary Figure S6

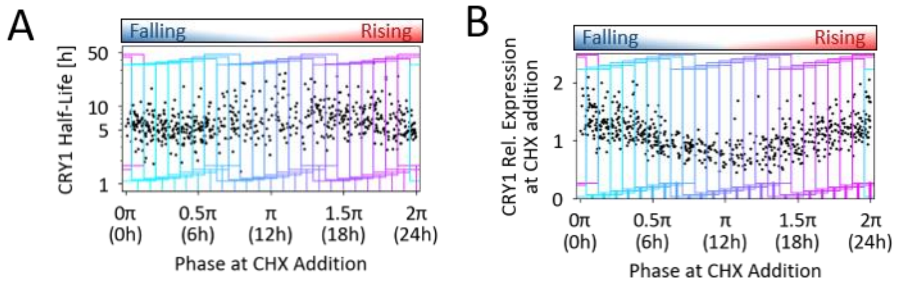


Supplementary Figure S5:

(A) Histogram of circadian phases of rhythmic cells at the time of CHX addition, 1-h bins, 10 cells per ring.

(B) Distribution of protein half-lives from cells transduced with control shRNA, with geometric mean and 95%CI. Data are pooled from several clones and 3 experiments. Half-lives longer than 30 h (< 0.5 % of all values) not shown.

(C) Distribution and median of protein half-lives from individual clones transduced with the indicated shRNA, data pooled from 3 experiments.

(D) Correlation of measured half-lives with circadian period length after knockdown of FBXL3. Correlation coefficient (r) and p-value from Spearman correlation, line: linear regression.

### Supplementary Figure S7

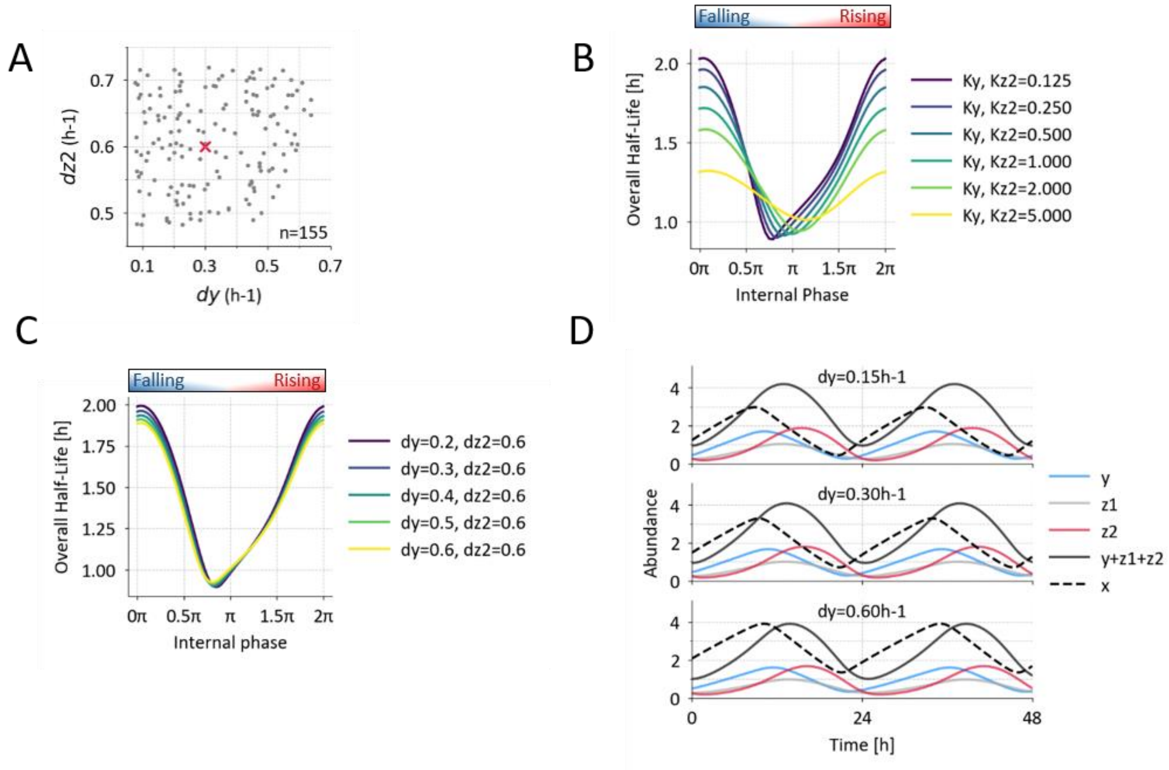


Supplementary Figure S6:

Example of binning: Single cell CRY1 protein half-life (A) or relative expression (B) values are binned into 24 overlapping sliding phase windows of ~0.78 rad (3h) for further analysis.

### Supplementary Figure S7

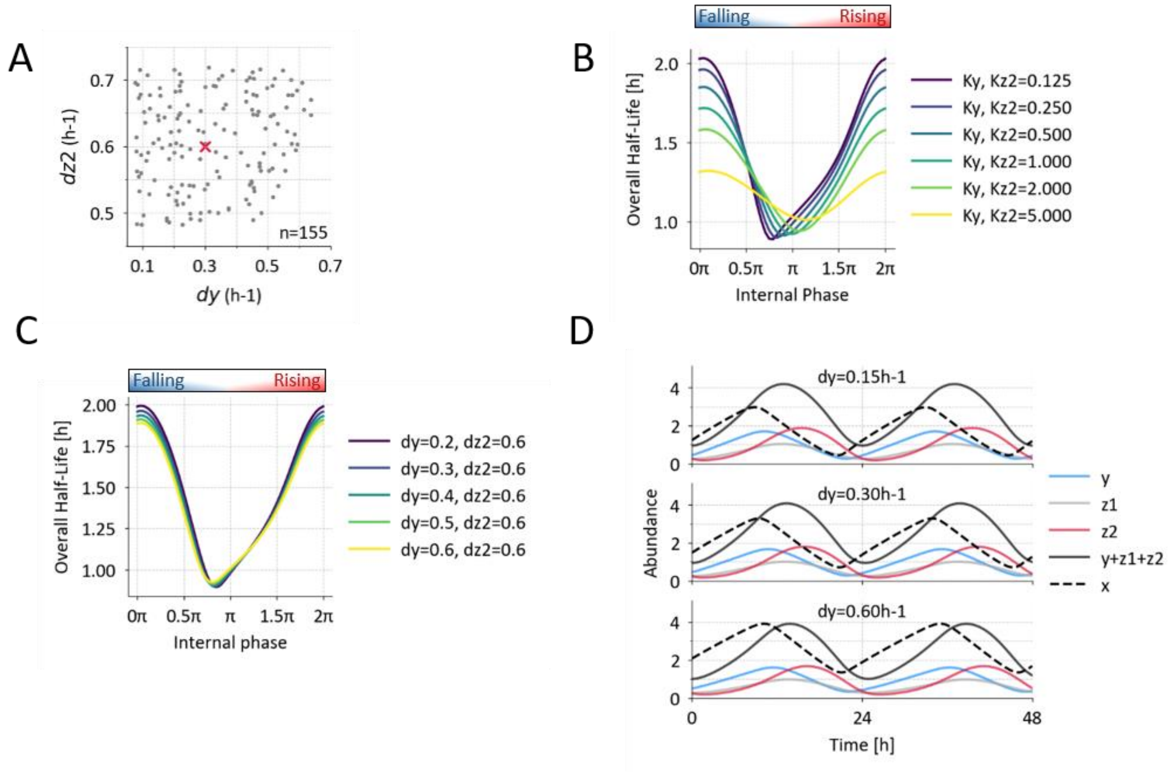


Supplementary Figure S7:

(A) Degradation rates of 'early' CRY1 (dy) and 'late' CRY1 (dz2) of the simulated population from Figure 5D-E, default values marked in red.

(B) Simulation of the phase-dependent half-life of the CRY1 pool for different rates of 'early' CRY1 degradation. The amplitude of stability decreases as degradation rates (dy and dz2) become more similar.

(C) Simulation of the phase-dependent CRY1 pool half-life for different $K_m$ values for the degradation of 'early' and 'late' CRY1. As rate limitation is relaxed by increasing $K_m$ values, the amplitude of stability is reduced.

(D) Simulation of CRY1 mRNA and subspecies abundance for different rates of 'early' CRY1 degradation. Lowest rate (highest stability) on top, default rate in middle panel.

### Supplementary Figure S8

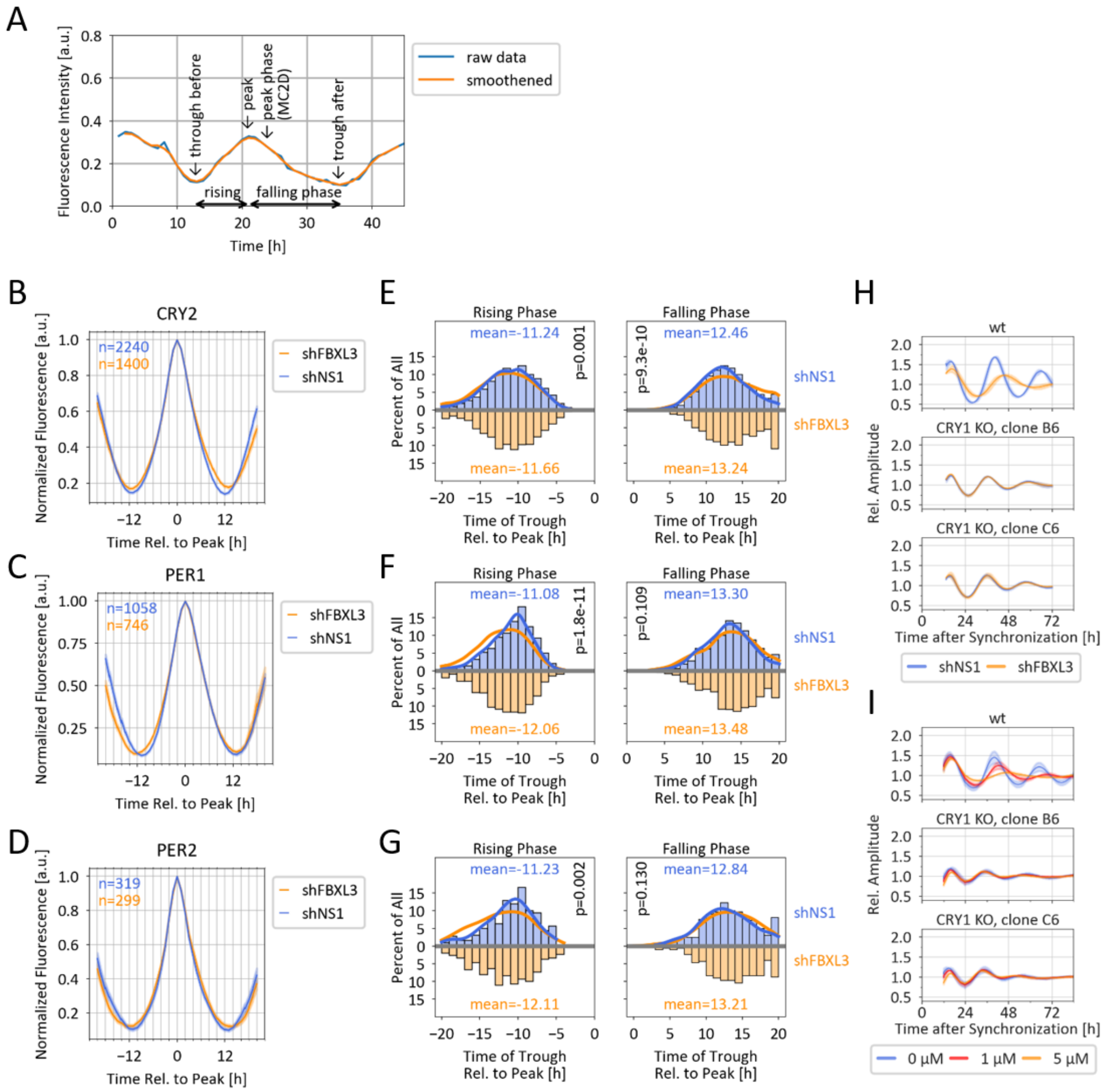


Supplementary Figure S8:

(A): Strategy to estimate trough-to-peak distance. Peak phase(MC2D): peak phase from Metacycle2D analysis.

(B-D) Average peak shape of CRY2, PER1 or PER2 expression after normalization (see Methods for details).

(E-G) Histogram of through-to-peak durations in the CRY2, PER1 or PER2 time series. Lines represent kernel density estimation. p-value: Mann-Whitney-U test, n as in (B-D).

(H-I) Detrended bioluminescence time series of wt and two *CRY1*$^{-/-}$ *Bmal1*:Luc reporter cell clones after dexamethasone synchronization. H: Cells were transduced with non-silencing or FBXL3-targeting shRNA. Mean + SD of n=2 experiments. I: Cells were treated with indicated concentration of KL001. Mean + SD of n=3 experiments.

### Supplementary Video Legends

#### Supplementary Video SV1:

Fluorescence image sequences of 25 nuclei per cell clone, acquired indicated hours after synchronization by dexamethasone.

#### Supplementary Video SV2:

Fluorescence image sequences of 25 nuclei per cell clone either treated with either solvent (top) or 20 µg/ml cycloheximide (CHX, bottom), acquired indicated hours before or after treatment.
